# DISCO-seq: 3D single-cell transcriptomics of intact biological systems

**DOI:** 10.64898/2025.12.10.693352

**Authors:** Harsharan Singh Bhatia, Laurent H.A. Simons, Louis B. Kuemmerle, Cristin McCabe, Sophie Jansen, Zhison He, Mayar Ali, Denise Jeridi, Mihail Ivilinov Todorov, Luciano Hoeher, Sudharsan Padmarasu, Ha Eun Park, Justus Flinn Thevis, Laura M. Bartos, Inderjeet Singh, Siegfried Ussar, Kıvanç Görgülü, Hana Algül, Kenny Roberts, Omer A. Bayraktar, Matthias Brendel, Fabian J. Theis, Barbara Treutlein, Aviv Regev, Ali Ertürk

## Abstract

Single-cell transcriptomics has transformed tissue analysis, yet current methods struggle to integrate whole-tissue 3D architecture. Conventional techniques restrict molecular profiling to pre-selected 2D sections, losing systemic context and introducing anatomical bias by sampling less than 0.001% of a whole organism. To overcome these challenges, we developed DISCO-seq, a tissue-clearing chemistry that enables superior RNA accessibility compared to fresh or fixed tissues. DISCO-seq integrates whole-organ or organism 3D imaging with both untargeted and targeted transcriptomics, yielding high-quality RNA from cleared tissues comparable to standard samples. We demonstrate its versatility by investigating tumor heterogeneity in a syngeneic glioblastoma mouse model, using 3D imaging to identify spatially distinct microenvironments and characterize their unique transcriptomic signatures. Moreover, DISCO-seq enabled unbiased, whole-body mapping of SARS-CoV-2 S1 protein deposition in mice, followed by transcriptomic profiling of spatially defined niches. By bridging mesoscale 3D imaging with single-cell transcriptomics, DISCO-seq establishes a paradigm for anatomically contextualized, hypothesis-free tissue interrogation.

**Graphical abstract:** 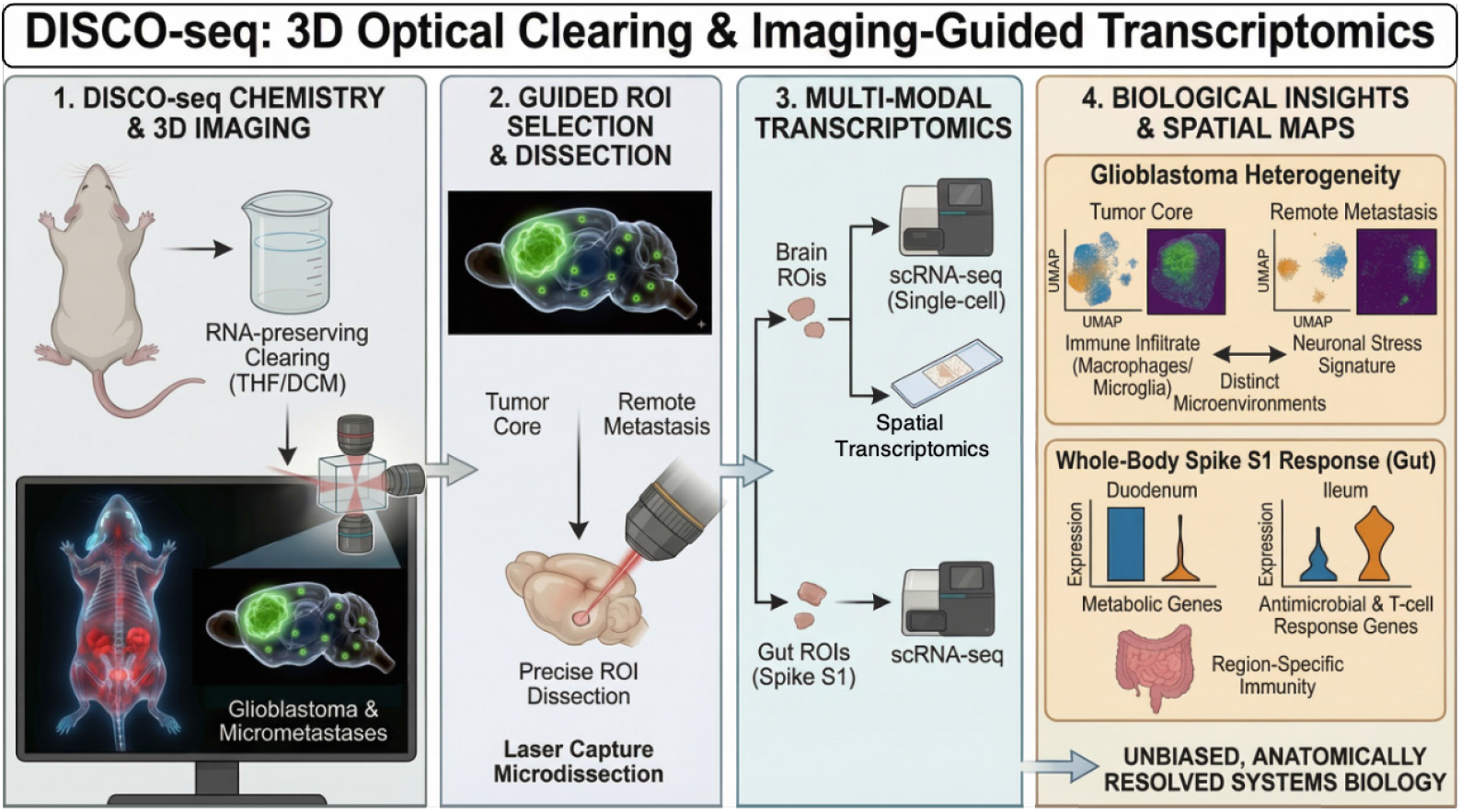

**Highlights:**

- DISCO-seq integrates RNA-preserving tissue-clearing chemistry with whole-organ or organism 3D imaging, enabling anatomically unbiased single-cell transcriptomics.
- DISCO-seq yields RNA quality and transcriptome profiles equivalent to those obtained from matched fresh or fixed tissues.
- DISCO-seq identifies discrete glioblastoma microenvironments and defines their transcriptomic states within the intact brain.
- DISCO-seq enables whole-body mapping of SARS-CoV-2 S1 and uncovers region-specific immune and metabolic responses across anatomical niches.

Supplementary movies can be seen at: http://discotechnologies.org/DISCO-seq/

## Introduction

Understanding gene expression within the spatial context of intact tissues and organs is fundamental to deciphering complex biological systems. Spatial transcriptomics technologies have enabled localization of RNA molecules within histological sections, offering unprecedented insights into cellular organization and tissue architecture (1–4). However, most existing platforms operate in two dimensions, relying on thin tissue sections. This limits the capacity to reconstruct complete organ- or organism-level molecular landscapes and is inadequate for applications requiring analysis of large tissue volumes or long-range cellular interactions, such as in neuroscience (5–10), infection biology (11, 12), and cancer pathology (13, 14). Furthermore, the common practice of selecting regions of interest (ROIs) from 2D sections introduces sampling bias and risks overlooking rare or pathologically relevant cell populations or microenvironments. This practice is particularly limiting at the organism level; for instance, a typical single-cell experiment profiling 100,000–200,000 cells from a mouse represents a vanishingly small fraction (∼0.001%) of the whole body. This introduces a profound sampling bias that hinders the unbiased discovery of rare pathological events, such as the initial seeding of a cancer cell. While methods like Expansion Microscopy (ExM) and its derivatives (15–17) or STARmap (18) offer high resolution or 3D capabilities, they face limitations in scalability, gene panel size, or compatibility with large-volume clearing, needed for whole-organ analysis. Other approaches based on in situ hybridization (ISH) or sequencing (3) (19) (20–23) are powerful but often require ROI preselection or struggle with the opacity and imaging depth constraints of large tissues.

The field of tissue clearing has rapidly expanded over the last decade to address high-resolution 3D imaging needs, enabling the visualization of specific cell populations in their native anatomical context across billions of cells. Dozens of methods employing hundreds of different chemicals, ranging from organic solvents, alcohols, to detergents and hydrogels, have been used to render biological tissues transparent. However, a major and persistent limitation of these techniques, particularly powerful solvent-based methods, is their harsh chemical nature, which typically leads to severe degradation of RNA. This fundamental incompatibility has largely precluded the integration of large tissue clearing with unbiased, whole transcriptome analysis, creating a critical bottleneck for spatial systems biology. The recent TRISCO method demonstrated the first whole volumetric RNA profiling, but it was restricted to only selected organs and a panel of targeted mRNA molecules (22). Therefore, we set out to address this challenge by developing a new chemistry specifically designed for robust RNA preservation for scalable, unbiased transcriptomics analysis of cleared tissues, including whole mouse bodies.

Through a systematic screen of various clearing chemical reagents and tissue preparation conditions, we identified a workflow that achieves excellent tissue transparency while maintaining high RNA quality (5, 8, 24). This effort resulted in DISCO-seq, a fully integrated, imaging-guided pipeline built on our novel RNA-preserving chemistry, which enables subsequent spatially resolved transcriptomic profiling on tissues previously considered incompatible with such analyses. By first rendering tissues transparent and imaging them in 3D at cellular resolution, DISCO-seq allows for the unbiased identification and selection of ROIs based on global anatomical context or specific biological features (e.g., disease lesions, cell distribution patterns) before molecular analysis. This decouples imaging from molecular workflows, preserving 3D anatomical integrity, providing high-quality RNA data, and enabling transcriptome-wide analysis across intact specimens without sectioning-induced bias. The method supports whole-specimen interrogation while retaining spatial and molecular resolution, offering a scalable framework for spatial systems biology.

Here, we demonstrate DISCO-seq’s versatility across multiple biological systems and transcriptomic platforms. We validate its performance in the mouse brain, showing high RNA quality and robust gene detection comparable to standard protocols. We apply it to reveal spatial transcriptomic heterogeneity in a mouse glioblastoma model guided by 3D tumor architecture and map systemic host responses to SARS-CoV-2 S1 protein deposition across the entire mouse body. Building on foundational advances in single-cell and spatial transcriptomics (25–28), DISCO-seq provides a physical framework for mapping cells and their molecular states within their native structural and functional contexts, aligning with the goals of initiatives like the Human Cell Atlas (29, 30).

## Results

### Optimization of RNA extraction and cDNA library preparation from cleared brain tissue

To establish a workflow for unbiased, three-dimensional spatial-molecular profiling, we first systematically addressed the critical challenge of RNA degradation during solvent-based clearing (6, 7, 31). We performed a comprehensive screen of tissue processing chemistries, evaluating several distinct con ditions that combined different perfusion reagents (PBS, RNAlater), fixatives (none, glyoxal, PFA), and clearing reagents (Tetrahydrofuran (THF)/ dichloromethane (DCM)/Benzyl-alcohol: Benzyl-benzoate (BABB; 1:2 ratio), Tertiary-Butanol (tert-butanol)/DCM/BABB) to measure their impact on RNA yield, purity, and integrity **(Table S1)**.

Our analysis revealed that while fixation-free clearing with RNAlater perfusion produced a high apparent RNA yield, it suffered from significant organic solvent contamination, rendering the RNA unsuitable for sequencing (A260/230 = 1.61-1.65). Conversely, alternative fixatives like glyoxal led to poor RNA pu ity (A260/280 = 1.72-1.91; A260/230 =0.7-0.9) and lower RNA integrity (RIN=2.3-2.4) **(Table S1)**. This systematic screen identified a precise workflow, the DISCO-seq chemistry, that robustly preserves RNA. The optimal protocol consists of cold PBS perfusion while maintaining the whole mouse on an ice bed, followed by 4% cold PFA fixation and optimized incubation with clearing reagents, i.e., THF and DCM. This combination yielded RNA of sequencing-grade quality (A260/280 = 2.10; A260/230 = 2.18; RIN=7-8.1), indistinguishable from fresh uncleared control samples **(Table S1)**. The success of this chemistry relies on a multi-step preservation strategy: initial PFA fixation cross-links the tissue and inactivates RNases; subsequent dehydration with a specific series of alcohols and organic solvents including THF and DCM, creates an anhydrous environment that further inhibits enzymatic activity; and finally, our optimized reverse-clearing and rehydration protocol gently removes hydrophobic residues to make RNA accessible for downstream enzymatic reactions. A careful transition back to aqueous conditions was necessary to prepare these cleared tissues for probe-based fixed RNA sequencing (10x Genomics Flex assay), which circumvents issues related to RT on damaged templates. We performed a gradual reverse clearing, first removing BABB with THF washes, followed by a stepwise rehydration using decreasing concentrations of THF (100, 90, 70, 50 %) in an aqueous buffer. This strategy aimed to gently reintroduce water, allowing fixed RNA molecules to regain a conformation amenable to probe hybridization while minimizing osmotic shock and guarding against any potential residual RNase activity during the transition. Once the cleared samples were fully rehydrated, either the samples were flash frozen at −80 degrees or immediately processed for RNA isolation and nano-drop-based measurement of RNA concentration **(Fig. 1A)**. We used fresh and fixed frozen brain samples as state-of-the-art control samples along with DISCO-seq cleared samples. As shown in **Fig. 1B**, total RNA was successfully extracted from fresh-frozen, fixed, and DISCO-seq cleared mouse brain tissues following optimized rehydration and crosslink reversal steps. RNA yield (range of 90-104 ng/µL) and purity (260/230 ratio ∼ 2.0) were comparable across conditions **(Fig. 1B)**, indicating minimal degradation and contamination of organic solvents in RNA extracted from cleared tissue. Next, we developed the isolation protocol of single cell suspension and single nucleus from fresh frozen and cleared brain tissues, which is described in detail in the method section **(Fig. 1C)**. Further, we validated cDNA library generation using the 10x Genomics Flex assay and SMART-seq2 from cleared single cells **(Fig. 1D)** and nuclei, respectively. The resulting electrophoretic profiles revealed consistent and specific library peaks around ∼262 bp in fresh and DISCO-seq cleared samples **(Fig. 1D)**, confirming successful amplification and integrity of the final sequencing libraries suitable for downstream single-cell RNA sequencing.

**Figure 1.**
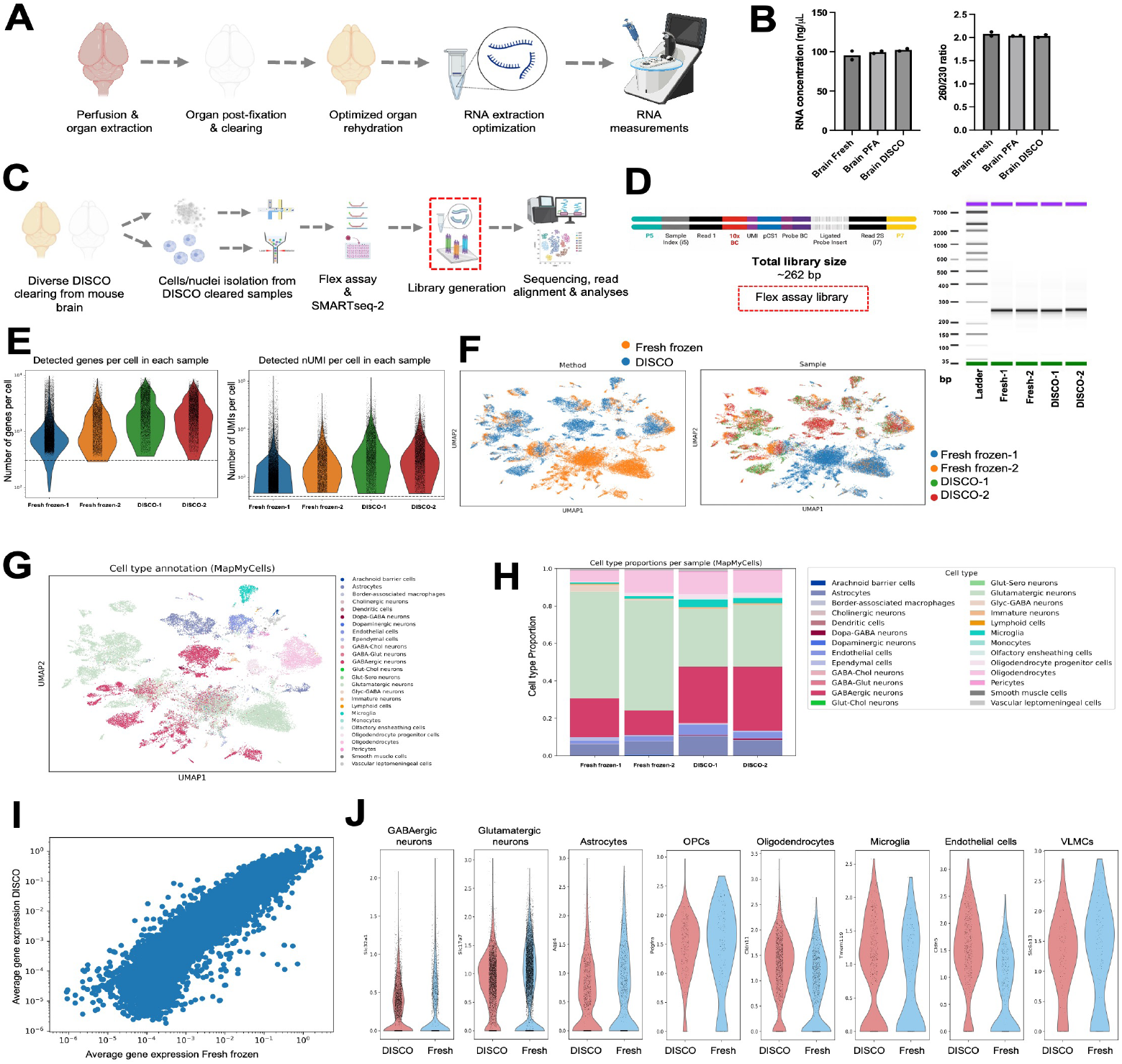
High-throughput single cell RNA sequencing in cleared mouse brains using DISCO-seq. **(A)** Schematic of DISCO-seq chemistry and RNA extraction optimizations from fresh frozen, PFA fixed and DISCO cleared brain samples. **(B)** Successful extraction of total RNA from fresh frozen, PFA fixed and cleared/rehydrated samples (left bar plot) with 260/230 values indicating purity of RNA (right bar plot). **(C)** Pipeline of cDNA library generation and sequencing from isolated single cells (10x Genomics Flex assay) and nuclei (SMART-seq2) from DISCO cleared samples. **(D)** Schematic showing the expected total library construct after the flex assay and electrophoretic gel showing the specfic bands of proper library construct of 262 bp in fresh and DISCO cleared samples. **(E)** Number of genes (left panel) and number of UMIs (right panel) recovered per cell in fresh frozen samples and DISCO cleared mouse brain samples. (N=2, total number of cells= 36,668, after filtering). **(F)** UMAP of the integrated scVI embedding colored by sample and **(G)** by annotated cell types. **(H)** Cell type proportions within each sample. **(I)** Scatterplots of average gene expression (Spearman correlation, R=0.94; Pearson correlation, R=0.80) between DISCO cleared samples vs. fresh frozen samples. **(J)** Representative gene expression of canonical marker genes in fresh frozen vs. DISCO samples.

### Scalable single-cell transcriptome profiling of cleared tissues

Following enzymatic dissociation and trituration, we obtained a single-cell suspension compatible with Flex chemistry. We compared fresh-frozen mouse brain samples with DISCO-cleared samples, analysed more than 36,668 cells (after filtering) with 20,000 mean reads per cell in an optimized protocol of 10x Genomics flex assay. Quality control of processed cells indicated that DISCO-seq cleared samples exhibited minimal doublet scores and maintained mitochondrial gene counts within 10 % of those observed in matched cell clusters from DISCO-seq vs. fresh samples **(Fig. S1A)**. DISCO-seq cleared samples exhibited higher numbers of detected genes per cell (median values of approximately 2,557 and 2,424 genes per cell for DISCO-seq sample 1 and 2, respectively) compared to control samples (median values of approximately 972 and 1,693 genes per cell for control sample 1 and 2, respectively), **(Fig. 1E)**. This suggests that the DISCO-seq clearing protocol may enhance RNA detection, possibly by improving tissue permeability and accessibility for sequencing reagents. Quantifying unique molecular identifiers (UMIs) per cell further supported the enhanced RNA detection in cleared samples. DISCO-seq cleared samples demonstrated significantly higher UMI counts (median values of approximately 3,672 and 3,380 UMIs per cell for DISCO-seq Sample 1 and 2, respectively) compared to control samples (median values of approximately 1,177 and 2,346 UMIs per cell for Control Sample 1 and 2, respectively) **(Fig. 1E)**. This substantial increase in UMI counts parallels the elevated gene detection rates, corroborating evidence that the THF/DCM-based clearing protocol enhances the detection of mRNA molecules in single-cell RNA sequencing. Unsupervised clustering and Uniform Manifold Approximation and Projection (UMAP) dimensionality reduction of high-quality cells revealed cellular populations distributed across conditions, regions and by cell types **(Fig. 1F-G, Fig. S1B)** with comparable cellular compositions across major annotated brain cell types **(Fig. 1H)**. Both sample types yielded high correlation of average gene expression (Spearman correlation, R=0.94; Pearson correlation, R=0.80) without significant difference of gene expression distribution for selected canonical marker genes **(Fig. 1I-J, Fig.S1C)**. This equivalence in cell-type representation and gene expression demonstrates that DISCO-seq preserves the cellular heterogeneity in native tissues, with minimal bias introduced by the clearing process.

### Transcriptome coverage in sorted nuclei from cleared tissues

Towards bulk and single-nucleus library preparation from diverse DISCO-clearing variants, we compared our DISCO-seq chemistry by assessing the impact of de-crosslinking and RNase inhibition on library quality **(See Table S1)**. As outlined in **Fig. S2A**, nuclei were isolated from cleared mouse brains, sorted, and processed for RNA and cDNA preparation. Bioanalyzer traces demonstrated higher cDNA integrity when de-crosslinking was performed at a certain temperature (see method section for optimization of protocol) prior to SMART-seq2 **(Fig. S2B–C)**, underscoring the importance of reversing crosslinks for efficient transcript recovery. Quantitative analysis across decreasing input numbers (500, 100, 50, 10, and single nucleus) revealed reproducible library size distributions for both DIS-CO-seq and tert-butanol based clearing method **(Fig. S2D–E)**. Inclusion of recombinant RNase inhibitor (RRI) further improved cDNA fragment size, particularly at higher input levels **(Fig. S2F–G)**. The protective effect of RNAse inhibition is particularly marked in the DISCO-seq protocol when compared with tert-butanol. Collectively, these developments establish a robust pipeline for generating high-quality bulk and single-nucleus cDNA libraries from chemically cleared tissues. To assess the transcriptome coverage and depth achievable with DISCO-seq, we performed RNA sequencing on both bulk and single-nucleus preparations from cleared mouse brain tissues. DISCO-seq cleared samples yielded slightly higher gene detection rates than tert-butanol cleared samples, particularly in bulk inputs **(Fig. S3A-B)**. With bulk preparations (500 nuclei), we detected around 12,000 genes in DISCO-seq. Down to the 10-nuclei level, DISCO-seq and tert-buta-nol-cleared samples consistently detected >4000 genes across RNase-inhibitor treated or untreated samples. Adding RNase inhibitors did improve the cDNA fragment size, however it did not further enhance the recovery of gene detection rates **(Fig. S3C-D)**, In the single nucleus experiment, both DISCO-seq and tert-butanol cleared samples showed >200 genes in a limited N of 17 out of 24 nuclei and 20 out of 24 nuclei, respectively (not separated based on RNase treatment) **(Fig. S3E-F)**. Importantly, among others, key cell-type marker genes for neuronal and glial populations were readily detectable in our datasets, including excitatory neuronal markers (*Satb2, Slc17a7*), inhibitory neuronal markers (Gad2, Meis2), and glial markers (*Sox10* for oligodendrocytes, and *Ptk2b* for microglia) **(Fig. S3G-H)**.

### Whole-brain volumetric mapping of glioblastoma dissemination using optical tissue clearing and 3D quantification

To showcase DISCO-seq’s utility in dissecting complex disease ecosystems, we applied it to a syngeneic mouse model of glioblastoma. SB28-eGFP cells were stereotactically injected into the left cerebral hemisphere of adult mice, and tumor progression was initially monitored via amino acid ([18F]FET) positron emission tomography (PET), which represents the clinical tool for molecular glioma imaging (32, 33), after three weeks following established protocols (34). While PET imaging provided valuable macroscopic information on tumor localization and growth dynamics, its inherent spatial resolution limitations (approximately 0.8-1.5 mm in dedicated small animal systems) precluded detailed cellular-level analysis of tumor heterogeneity and microenvironmental interactions even at a stage of large tumors. In contrast, DISCO-seq, when combined with whole-brain optical tissue clearing, enabled comprehensive single-cell resolution mapping of tumor cells and their surrounding neural microenvironment across the entire brain, offering unprecedented insights into the spatial distribution and molecular characteristics of the glioblastoma ecosystem **(Fig. 2A)**. Following whole-brain clearing and 3D imaging, we observed a large primary tumor mass with several discrete satellite regions of tumor cell accumulation and their trafficking to contralateral region via corpus callosum **(Fig. 2B, Video S1)** which were not visible in amino acid PET imaging in week 3. We further developed a novel method to co-register and quantify these PET/CT and light sheet scans of glioblastoma brain with the Allen brain atlas, revealing all the tumors, including single disseminated ones, in a whole-brain region-specific manner **(Fig. 2C-D, Table S2)**. Whole-brain volumetric analysis of tumor burden revealed a strikingly asymmetric distribution of lesions across the primary (38 annotated) brain regions. Quantitative segmentation of the optically cleared and 3D-imaged whole brain delineated three categories of tumor architecture: small tumor blobs, tumor clusters, and a large contiguous tumor. Across all regions, the large tumor mass contributed the vast majority of the total tumor volume (3.57 × 10^10^ voxels; ∼99% of the overall burden), while small and clustered lesions accounted for only 9.7 × 10^7^ voxels (0.27%) and 2.19 × 10^8^ voxels (0.61%), respectively. The smaller lesions were widespread and biologically significant despite their limited volumetric fraction. The largest contiguous tumor spanned multiple anatomically and functionally distinct territories, including the hippocampal formation, thalamic nuclei, somatosensory cortex, cerebral nuclei, fiber tracts, and midbrain motor regions, with local volumes ranging from 4 × 10^8^ to 1.8 × 10^10^ voxels per region. This pattern indicates highly invasive growth along white-matter pathways and perivascular routes, consistent with the known tropism of glioblastoma for myelinated tracts and vascular niches.

**Figure 2.**
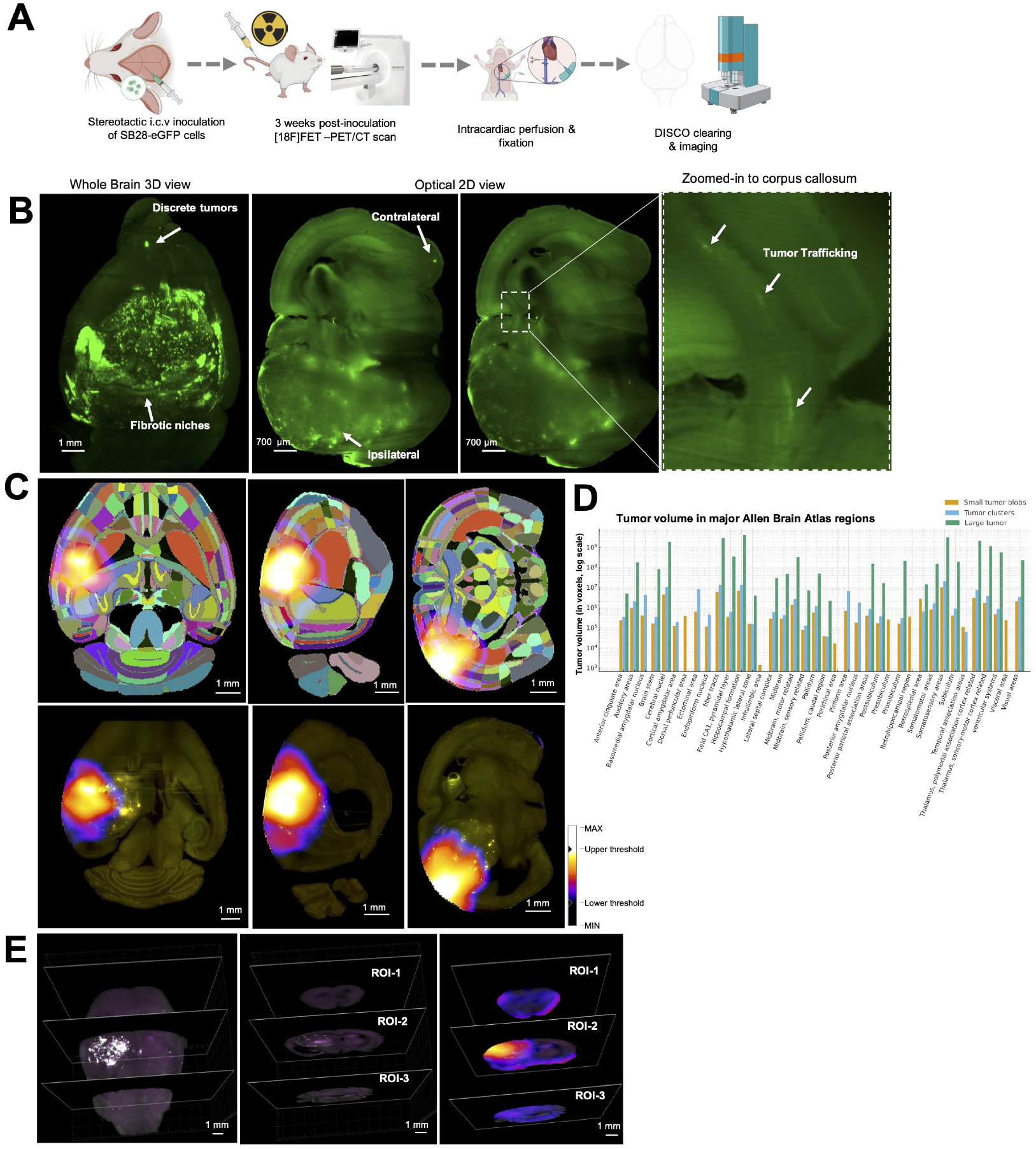
3D imaging guided detection of single discrete tumor cells and their trafficking in syngeneic mouse model of Glioblastoma. **(A)** Experimental pipeline of mouse model of Glioblastoma (syngeneic mouse model of SB28-eGFP injected with tumor cells). **(B)** 3D Images of whole brain showing large primary tumor and few discrete regions associated with small/single or aggregated tumor cells (channel 488 labeled in green tumor cells) and their trafficking via corpus callosum (N=7 glioblastoma, sham, N=4). **(C)** PET-CT-lightsheet co-alignment of glioblastoma whole brains using hierarchically and randomly color-coded Allen brain atlas to reveal annotated regions. 32-bit scale indicates min-max range of 11-1630 with lower threshold of 373 to higher threshold 1244. **(D)** Quantification of virtual reality (VR) segmented tumor volume in major regions of whole glioblastoma brain. **(E)** 3D images of whole brain imaged with 4x objective showing big tumor in white (488 channel), magenta is autofluorescence (594 channel) with 3-ROIs selection from hot vs. cold tumor regions in whole brain.

In contrast, the small tumor blobs and tumor clusters were observed as discrete, spatially dispersed foci throughout cortical and subcortical areas - including the visual cortex (2.0 × 10^6^ voxels), posterior parietal association areas (3.9 × 10^5^ voxels), and entorhinal and perirhinal cortices (6.5 × 10^5^ voxels combined) - as well as in deep structures such as the cerebral nuclei (4.6 × 10^6^ voxels) and fiber tracts (6.0 × 10^6^ voxels). Although volumetrically small, these microlesions were widely disseminated across the brain, often separated by millimeter-scale distances, and suggest metastatic or migratory seeding events that would be invisible to conventional imaging modalities.

Next, we defined specific fibrotic niches characterized by dense cellular aggregation at the injection site, and infiltrative regions (rostral and caudal regions) showing dispersed tumor cells spreading potentially via unknown molecular mechanisms **(Fig. 2E)**.

### DISCO-seq enables spatially-resolved transcriptomic profiling of glioblastoma infiltration patterns

We combined optical tissue-clearing-guided regional dissection with high-throughput single-cell RNA sequencing to uncover the molecular heterogeneity of disseminated glioblastoma cells at a whole-brain scale. Imaging of cleared glioblastoma brains identified a primary core tumor region (ROI-2) and two anatomically remote regions (ROI-1 and ROI-3) containing sparse, GFP^+^ tumor cells that had potentially migrated across the corpus callosum from the inoculation site **(Fig. 3A)**. These precisely localized ROIs were subjected to high-throughput scRNA-seq together with matched regions from sham-injected control brains, thereby enabling direct region- and condition-specific transcriptomic comparison. Following single-cell RNA sequencing, we recovered high-quality transcriptomes that maintained the expected cellular diversity and proportions characteristic of brain tissue. Quality control metrics confirmed the robust preservation of RNA integrity across all samples processed through the DISCO-seq clearing protocol. Doublet scores remained consistently low across all conditions and cell types, validating the quality of cellular dissociation and downstream processing **(Fig. S4A)**. Mitochondrial gene count percentages were comparable between sham and glioblastoma samples across all three ROIs, indicating minimal stress-induced artifacts from the clearing procedure **(Fig. S4B)**.

**Figure 3.**
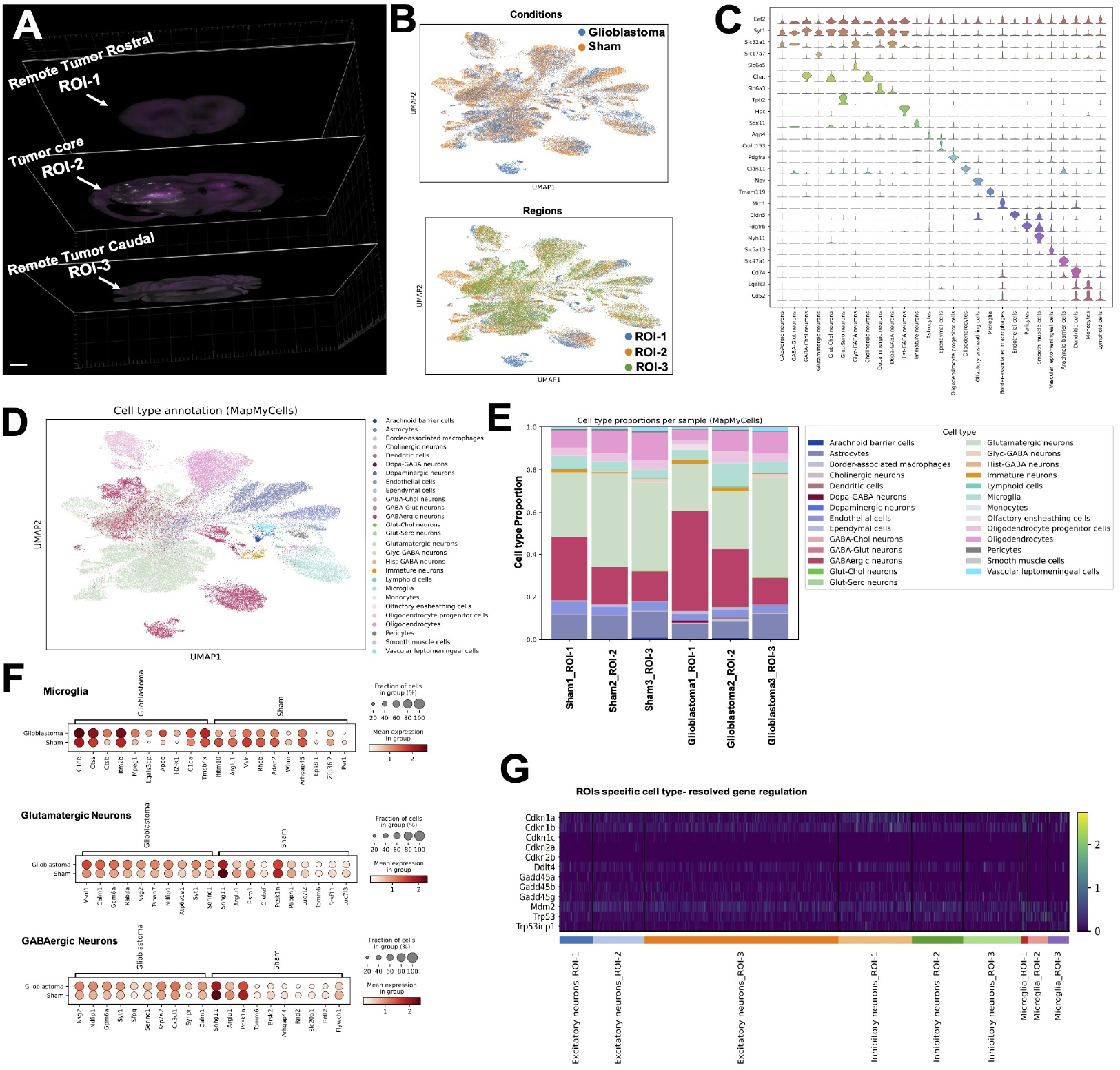
DISCO-seq detects regional transcriptomic signatures of disseminated tumor cells in Glioblastoma mouse model. **(A)** Whole glioblastoma brain showing the primary core tumor region (ROI-2) and remote regions (ROI-1, ROI-3) as shown in Fig. 2E. **(B)** UMAPs showing the distribution of cells per conditions (upper panel) and per regions (lower panel), N=3 per condition, 3 ROIs pooled per sample, total number of cells= 62,782. **(C)** Violin plots showing the expression of canonical marker gene expression across cell types with representative housekeeping marker gene (Eef2). **(D)** UMAP colored by cell type annotations. **(E)** Cell type proportions of each sample. **(F)** Top ten differentially expressed genes (DEG) in microglia/myeloid cells (upper dot plot), in glutamatergic neurons (middle dot plot), and in GABAergic neurons (lower dot plot) in glioblastoma compared to sham brains. **(G)** ROIs dependent (roastral to caudal) neuronal sub-types (excitatory and inhibitory) and microglia specific regulation of specific stress-related and DNA damage related genes.

We recovered a median of 2,129 genes per cell with approximately 2,823 UMI counts per cell, demonstrating sufficient sequencing depth for reliable cell type identification and differential expression analysis **(Fig. S4C)**. Notably, these metrics showed remarkable consistency across sham controls (Sham1_ ROI-1, Sham2_ROI-2, Sham3_ROI-3) and their corresponding glioblastoma regions (Glioblastoma1_ ROI-1, Glioblastoma2_ROI-2, Glioblastoma3_ROI-3). Unsupervised clustering and UMAP dimensionality reduction of 62,782 high-quality cells revealed distinct cellular populations distributed across conditions and regions **(Fig. 3B)**. The upper UMAP in **Fig. 3B** demonstrated clear separation between sham and glioblastoma samples. In contrast, the lower panel in **Fig. 3B** revealed region-specific clustering patterns corresponding to the three anatomical ROIs. We identified 27 major cell types based on canonical marker gene expression, including astrocytes, oligodendrocytes, oligodendrocyte precursor cells (OPCs), microglia, macrophages, neurons (subdivided into GABAergic, glutamatergic, dopaminergic, cholinergic, immature subtypes), ependymal cells, endothelial cells, pericytes, and vascular smooth muscle cells **(Fig. 3C-D)**. Expression of the housekeeping gene *Eef2* was uniformly distributed across all cell types, confirming technical consistency **(Fig. 3C)**. The comprehensive marker gene analysis, including cell-type-specific markers with their corresponding mean expression levels and fraction of expressing cells, validated our annotation strategy **(Fig. S4D-E)**. To evaluate the transcriptional diversity and coverage of the DISCO-seq dataset, we quantified the number of genes detected per annotated cell type **(Fig. S4F)**. Across the entire dataset, the analysis revealed robust capture of significant neural, glial, and immune populations, with a detection range of 13k-17k genes per cell type across most classes. The numbers above each bar indicate the total number of cells contributing to each cluster, highlighting the depth and breadth of sampling across brain-resident populations. Notably, the uniform distribution of detected genes across diverse cell types-including low-frequency populations such as monocytes (20 cells), dendritic cells (109 cells), and diverse sub-sets of neuronal population-demonstrates the technical robustness of the optical tissue clearing and single-cell RNA-seq integration. These results confirm that DISCO-seq preserves transcriptional complexity across all major cell types, enabling quantitative comparison of both abundant and rare populations in the glioblastoma and control brains Quantitative analysis of cell type proportions revealed profound alterations in the cellular composition of the tumor microenvironment **(Fig. 3E)**. The most striking observation was a marked expansion of microglia/ macrophages in Glioblastoma2_ROI-2 (tumor core) compared to Sham2_ROI-2, increasing from approximately ∼5% to ∼12% of total cells. This three-fold enrichment was accompanied by differential expression of established glioblastoma-associated genes such as *C1qa, Ctss, Ctsb, Im2b, Mpeg1* in the myeloid compartment **(Fig. 3F)**. Global transcriptomic comparison between glioblastoma and sham conditions revealed pronounced neuronal subtype–specific reprogramming. Both glutamatergic and GABAergic neurons in glioblastoma exhibited coordinated upregulation of genes associated with synaptic signaling and vesicle trafficking (*Syt1, Rab3a, Gpm6a, Nsg2, Calm1, Nd-fip1*), indicative of enhanced excitability and neuronal remodeling. In contrast, transcripts linked to neuropeptide processing, stress regulation, and homeostatic balance (*Snhg11, Pcsk1n, Arhgap44*) were consistently downregulated, suggesting a loss of inhibitory control and altered neurosecretory function **(Fig. 3F)**. These global changes imply a shift toward hyperactive, tumor-interactive neuronal states that may facilitate glioma–neuron coupling. Neuronal population showed substantial depletion in the tumor core, decreasing from approximately 65% in sham tissue to 56% in the corresponding glioblastoma region. Astrocytes and oligodendrocyte lineage cells maintained relatively stable proportions across conditions, though subtle shifts were observed. Notably, the remote regions (ROI-1 and ROI-3) displayed intermediate phenotypes, with cellular proportions falling between sham controls and the tumor core, suggesting progressive microenvironmental remodeling extending beyond the primary tumor mass. To explore the spatial dimension of these effects, ROI–resolved analysis revealed substantial heterogeneity in the expression of stress- and cell cycle–associated genes across neuronal and glial populations. Genes within the p53–DNA damage–stress axis (*Cdkn1a/b/c, Gadd45* family, *Ddit4, Mdm2, Trp53, Trp53inp1*) displayed region-specific activation patterns, varying among excitatory, inhibitory, and microglial compartments. This ROI-dependent modulation underscores the localized nature of glioblastoma-induced transcriptional remodeling, where neuronal hyperactivity in tumor-adjacent regions is accompanied by selective stress and checkpoint signaling within both neurons and microglia. **(Fig. 3G, Table S3-S6)**. Collectively, these spatially-resolved single-cell transcriptomic analyses demonstrate that DISCO-seq successfully captures the complex cellular ecosystem of glioblastoma, revealing both the dramatic remodeling of the tumor core microenvironment and the subtle but significant molecular signatures of tumor infiltration in anatomically remote brain regions.

### Cross-validation of DISCO-seq data with single-cell resolution spatial transcriptomics in cleared samples

The above transcriptomic findings were further corroborated with high-resolution spatial transcriptomics using the Xenium In Situ platform (10x Genomics) on DISCO-seq cleared and imaged brain tissues. The Xenium platform enables high-resolution spatial transcriptomics by detecting RNA molecules directly within intact tissue sections using barcoded probes and iterative fluorescence imaging. Utilizing the Xenium mouse gene expression panel, we achieved robust signal detection and transcript accuracy, demonstrating the compatibility of advanced spatial transcriptomics with DISCO-seq tissue clearing chemistry. This enabled subcellular-resolution mapping of hundreds of transcripts from 3D navigated tumor remote environments. To this end, 3D whole brain DISCO clearing and imaging guided the ROI selection for spatial transcriptomics. The spatial distribution of identified cell types corresponded well with the expected brain region-specific patterns **(Fig. S5A)**. Both coarse and fine cell-type annotations revealed diverse cellular identities with regional specificity **(Fig. S5B-D)**, such as GABAergic olfactory bulb immature neurons (OB-IMN-GABA), which aligned well with published MERFISH brain atlas data (35). Importantly, DISCO-seq clearing enabled navigation and selection of remote regions throughout the whole brain that would remain undetectable without prior 3D visualization. This multi-platform compatibility highlights DISCO-seq’s versatility across long-read sequencing (SMART-seq2), high-throughput targeted whole transcriptome sequencing, and spatial subcellular-resolution RNA mapping via in situ techniques (Xenium). To evaluate the concordance between single-cell and spatial transcriptomic modalities, we compared DISCO-seq and spatial Xenium datasets derived from glioblastoma tissue. High-confidence cell-type assignments were first obtained using mapmycells, retaining only cells with mapping confidence ≥ 0.8 and excluding cell types represented by fewer than 80 cells in either dataset. This ensured that only robust and well-represented populations were included for downstream analysis. A dotplot was generated to visulaize the mean normalised gene expression (color intensity) and fraction of expressing cells (dot size) across matched cell-type groups in both datasets. The analysis revealed a striking consistency between DISCO-seq (red) and spatial Xenium (blue) profiles **(Fig. S5E)**, with canonical markers such as *Gfap* and *Aqp4* in astrocytes, *Pdgfra* and *Opalin* in oligodendrocyte lineage cells, and *Pecam1* and *Cldn5* in endothelial cells-showing concordant expression patterns across modalities. These results confirm that the spatial transcriptomic mapping faithfully reproduces transcriptional signatures captured by single-cell sequencing. To further quantify the agreement between modalities, we computed the average gene expression per cell type across all shared genes and compared them using Pearson correlation. The resulting scatterplot showed a strong positive correlation (R = 0.81) between DISCO-seq glioblastoma and spatial Xenium glioblastoma datasets **(Fig. S5F)**, demonstrating high reproducibility and alignment of molecular profiles at both the cell-type and gene-expression levels. Together, these analyses validate the integration of DISCO-seq and Xenium platforms for high-resolution spatially anchored single-cell transcriptomics in glioblastoma.

### DISCO-seq enables whole-body investigation of spike S1 distribution and regional transcriptome profiling in mouse small intestine

To demonstrate DISCO-seq’s capability for whole-organism analysis, we investigated the systemic distribution and host response to SARS-CoV-2 spike protein S1 in wild-type mice. Thirty minutes after intravenous administration of fluorescently labeled S1 protein (Alexa Fluor 647), we performed whole-body clearing and light-sheet imaging to map S1 distribution across multiple organs and found earlier known (11) as well as less characterize organs **(Fig. 4A)**. This unbiased approach revealed organ-specific tropism patterns, with pronounced S1 accumulation observed in the expected organs liver, lung, intestine, kidney with relatively sparse accumulation in heart, skeletal muscle and bone marrow (data not shown). High-resolution imaging of the small intestine revealed distinct S1 protein localization patterns **(Fig. 4B)**, with notable regional heterogeneity between the duodenum and ileum **(Fig. 4C-D, Video S2)**. Following gut extraction **(Fig. 4E)**, we dissected specific regions of the duodenum and ileum for down-stream single-cell RNA sequencing analysis **(Fig. 4F)**. Single-cell RNA sequencing of dissected duodenum and ileum regions yielded high-quality cellular transcriptomes across both PBS control and spike S1-treated conditions. Quality control metrics demonstrated robust data quality, with low doublet scores across all conditions and cell types **(Fig. S6A)**, minimal mitochondrial contamination **(Fig. S6B)**, and consistent recovery of genes and UMIs per cell **(Fig. S6C-D)**. UMAP dimensionality reduction revealed subtle, distinct cellular clustering patterns between spike S1-treated and PBS control samples **(Fig S6E, Fig. 4G)**, as well as clear regional segregation between duodenum and ileum-derived cells **(Fig. 4H)**.

**Figure 4.**
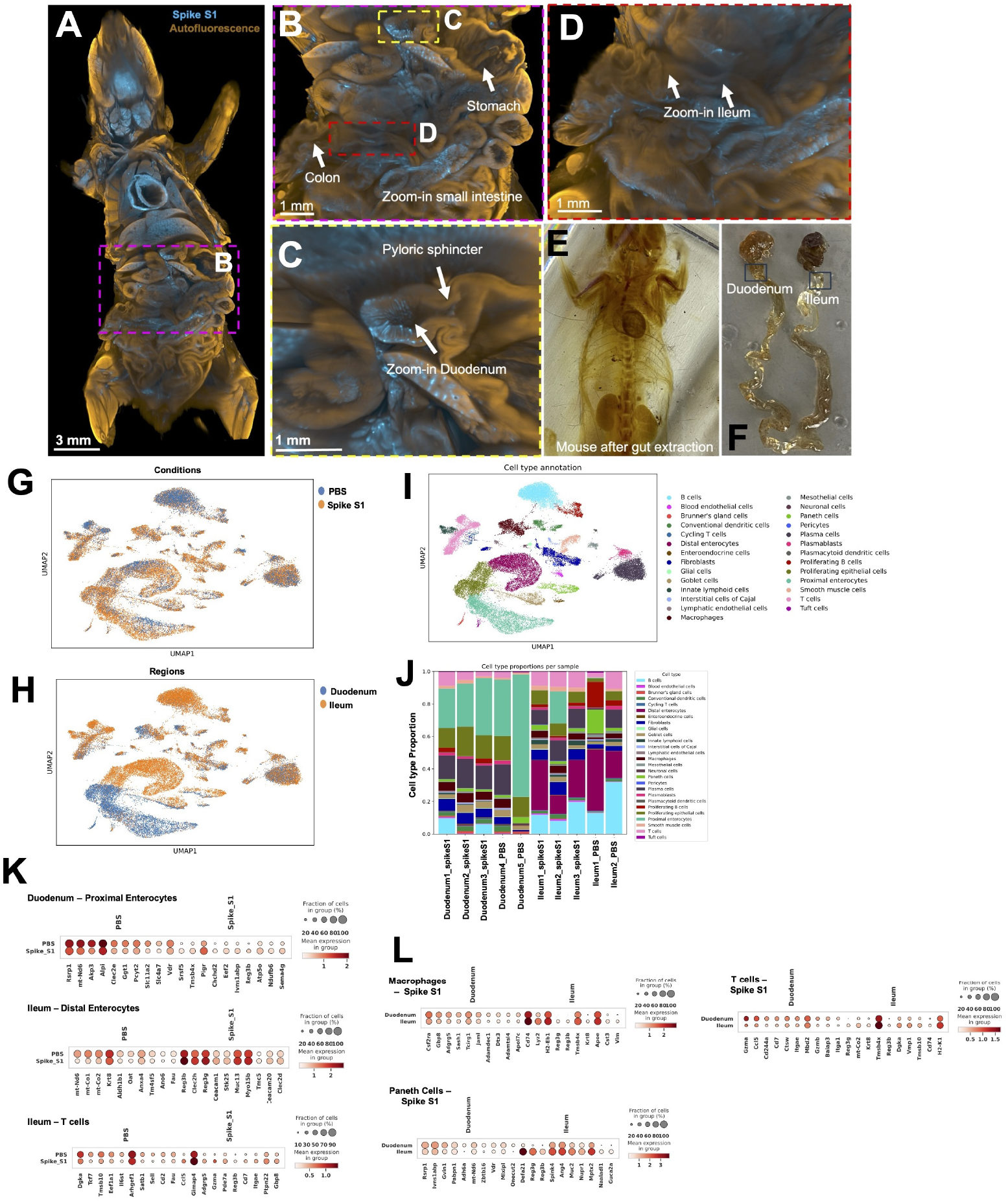
DISCO-seq enables whole body investigation of spike S1 distribution in mice and sub-regional transcriptome analyses of small intestine. **(A-F)** Unbiased investigation of spike S1 protein tropism in whole body of wild type mice after 30 min injection of spike S1 conjugated with Alexa fluorophore 647 (N=3 Spike S1; N=2 PBS controls, number of total cells= 37,853). **(B)** Zoomed-in small intestine. **(C)** Spike S1 tropism in duodenum. **(D)** Spike S1 tropism in ileum. **(E)** Whole cleared mouse after extraction of gut. **(F)** Regions of duodenum and ileum used for downstream scRNA processing. **(G)** UMAP of the small intestine’s scRNA-seq data colored by condition, **(H)** by region. **(I)** by cell type. **(J)** The relative cell proportion distribution of cell type within each condition. **(K)** Top ten differentially expressed genes (DEG) in proximal enterocytes (upper plot), distal enterocytes (middle plot) and in T cells from Ileum (bottom left plot). **(L)** Top ten differentially expressed genes (DEG) in region specific manner in macrophages (top left plot), Paneth cells (bottom left plot) and in T cells (top right plot) after spike S1 injection.

Comprehensive cell-type annotation identified major intestinal populations, including epithelial cells, T cells, B cells, plasma cells, stromal cells, and myeloid cells **(Fig. 4I)**. Cell-type identities were validated using canonical marker genes, which showed expected expression patterns and cellular fractions **(Fig. S6F)**. The relative proportions of cell types were generally consistent across conditions **(Fig. 4J)**, indicating that acute S1 exposure did not dramatically alter the overall cellular composition within the 30-minute timeframe examined. Differential gene expression analysis revealed that spike S1 protein elicited distinct transcriptional responses across multiple cell types, with notable regional differences between the duodenum and the ileum **(Table S7-S9)**. In the duodenum, proximal enterocytes upregulated genes involved in mitochondrial respiration and oxidative phosphorylation (*mt-Nd6, mt-Co1, Akp3, Ggt1, Pcyt2*), reflecting enhanced metabolic activation. In contrast, distal enterocytes showed induction of stress- and immune-interaction pathways (*Krt8, Aldh1b1, Clec2h, Reg3g*), suggesting barrier adaptation and local immune crosstalk. In the ileum, Paneth cells exhibited strong activation of antimicrobial and secretory programs (*Reg3g, Reg3b, Defa5, Ang4*), while macrophages upregulated inflammatory and antigen-presentation genes (*Cd74, Lyz2, Apoe, Tmsb4x*), indicating regionally enhanced mucosal immunity. T cells in the ileum showed coordinated expression of activation and cytotoxicity markers (*Gzma, Cd44, Itga4, H2-K1*), pointing to increased immune surveillance compared with the duodenum **(Fig. 4K-L)**.

Gene ontology enrichment supported these cell-type– specific transcriptional signatures: duodenal enterocytes predominantly activated mitochondrial and ATP metabolic processes, whereas ileal epithelial and immune cells engaged defense, antimicrobial, and cytokine response pathways **(S6G-H)**. These findings demonstrate that DISCO-seq captures subtle spatial heterogeneity in immune cell responses within anatomically adjacent segments of the intestinal tract. The regional specialization of T cell transcriptional programs-evident within just 30 minutes of systemic S1 exposure-suggests that local microenvironmental cues, tissue-resident cellular networks, or regional differences in S1 bio-availability critically shape immune cell function. The ability to resolve such fine-scale spatial biology represents a key advantage of the DISCO-seq approach, enabling the discovery of region-specific molecular responses that would be masked by conventional bulk tissue analysis. Together, these results demonstrate that DISCO-seq provides high-quality, biologically meaningful transcriptomic data from optically cleared specimens, with performance comparable to standard methods while offering the unique advantage of three-dimensional anatomical context and unbiased, imaging-guided molecular profiling. The method’s versatility across multiple RNA technologies-from untargeted long-read sequencing to high-throughput droplet-based methods and *in situ* hybridization approaches-makes it a powerful tool for comprehensive spatial systems biology in both research and clinical settings.

## Discussion

DISCO-seq represents a paradigm shift in spatial transcriptomics by enabling unbiased, imaging-guided identification of tissue regions for molecular profiling with preserved three-dimensional anatomical context. By integrating whole-organism optical tissue clearing with untargeted transcriptomics, DISCO-seq overcomes critical limitations of conventional section-based methods, including sampling bias, loss of global context, and limited scalability to whole organs or organisms.

### Technical innovations and advantages

The spatial transcriptomics field has evolved from pioneering section-based methods (1, 2) to high-resolution platforms such as MERFISH (4, 19) and three-dimensional approaches including STARmap (18), EA-SI-FISH (21), and expansion sequencing (17). However, these methods either rely on thin sections necessitating region pre-selection, or face constraints in gene coverage and tissue penetration depth. The TRISCO method pioneered the profiling of specific mRNAs in cleared, intact tissues; however, its reliance on probe-based *in situ* hybridization restricts its application to a pre-defined panel of targeted genes. DISCO-seq fundamentally overcomes these limitations by enabling, for the first time, untargeted, whole-transcriptome sequencing directly from these cleared volumes. Unlike recent probe-based clearing methods (22, 23) that depend on predetermined gene panels and require anatomical coordinate pre-selection, DISCO-seq employs imaging-guided dissection followed by untargeted RNA sequencing. This preserves RNA integrity while enabling capture of complete transcriptomes. By rendering entire organs transparent prior to region selection (5, 7, 31), DISCO-seq eliminates anatomical bias, revealing invisible microlesions, rare populations, or dispersed pathological features. Our validation demonstrates that THF/DCM-based cleared tissues yield data quality exceeding fresh-frozen samples, with enhanced gene detection (median ∼3,600 UMIs/cell) likely reflecting a key advantage of the clearing process itself: improved tissue permeability, which enhances probe accessibility and leads to more complete transcript capture. Critically, DISCO-seq maintains compatibility with multiple platforms-droplet-based methods, SMART-seq2 (28), and spatial transcriptomics-providing unprecedented flexibility across biological scales.

### Biological insights from glioblastoma and spike protein studies

Our glioblastoma analysis revealed extensive micrometastatic dissemination comprising only 0.88% of tumor volume yet distributed across multiple brain regions-invisible to clinical imaging. Spatially resolved profiling identified profound molecular heterogeneity extending beyond visible margins, with the tumor core showing dramatic immune remodeling (microglia/ macrophages expanding from 5% to 14%) while infiltrative regions displayed intermediate phenotypes with neuronal stress signatures. These findings align with recent whole-tumor studies (36, 37) demonstrating spatial evolutionary trajectories critical for therapeutic resistance. This approach holds particular promise for addressing the critical clinical challenge of recurrence prediction in glioblastoma patients; with larger patient cohorts, DISCO-seq-derived spatial molecular maps could be correlated with longitudinal recurrence patterns to generate predictive models for patient-specific relapse risk. The immune heterogeneity carries particular significance for immunotherapy development. The “cold” immune phenotype in tumor cores, characterized by immunosuppressive markers, contrasts sharply with distinct transcriptional states in infiltrative regions (38, 39), suggesting spatially variable therapeutic responses that bulk sampling would obscure. Whole-body spike S1 protein mapping revealed organ-specific tropism extending beyond ACE2 expression predictions (40). The striking regional T cell response dichotomy- duodenal T cells prioritizing transcriptional reprogramming versus ileal T cells activating robust antimicrobial defense-reveals previously unappreciated spatial immune specialization reflecting distinct microenvironmental cues, microbiome composition, and lymphoid organization (41, 42). The rapid responses (30 minutes) indicate spike protein alone triggers inflammatory cascades potentially contributing to post-acute COVID-19 sequelae (11), suggesting therapeutic interventions must account for organ-specific immune states.

### Implications and future directions

DISCO-seq exemplifies a conceptual shift from analyzing “where genes are expressed” to understanding “what molecular programs define anatomically observed niches.” This imaging-to-molecules workflow aligns with Human Cell Atlas goals (29, 30), avoiding computational artifacts from reconstructing three-dimensional models from sections while providing anatomically faithful mapping across species and disease states. While achieving cellular resolution across intact specimens, methodological considerations remain. Organic solvent clearing may affect highly fibrotic or calcified regions, requiring careful quality control for pathological specimens. The method currently relies on regional dissection rather than true volumetric *in situ* transcriptomics-integration of unbiased sequencing compatible with intact animal would enable direct spatial profiling. Future iterations integrating expansion microscopy (37) or lineage barcoding (43) could link transcriptional states to subcellular organization and cellular histories. DISCO-seq’s applications extend beyond our demonstrations. In developmental biology, it can enable spatiotemporal mapping across embryos; in neuroscience, brain-wide circuit mapping; in immunology, whole-lymphoid-organ response dynamics; and in precision oncology, comprehensive molecular tumor resection margin characterization for patients follow-up and risk stratification. The method’s whole-body mapping capacity has particular relevance for systemic diseases, including metastatic cancer (14), autoimmune disorders, and chronic infections.

## Conclusion

DISCO-seq provides a robust framework for unbiased spatial transcriptomics in intact specimens, bridging scales from whole organisms to single cells. By preserving complete anatomical context while enabling cellular-resolution molecular profiling, DISCO-seq addresses fundamental limitations of conventional approaches. As spatial biology transitions from describing locations to understanding how spatial organization shapes molecular states, integrated approaches coupling three-dimensional visualization with comprehensive molecular characterization will be essential for advancing our understanding of tissue organization, disease pathogenesis, and therapeutic responses across basic research and clinical diagnostics.

## Acknowledgements

This work was supported by the Vascular Dementia Research Foundation, Deutsche Forschungsgemeinschaft (DFG, German Research Foundation) under Germany’s Excellence Strategy within the framework of the Munich Cluster for Systems Neurology (EXC 2145 SyNergy, ID 390857198), ERC Consolidator Grant (AE, GA 865323), Nomis Heart Atlas Project Grant (Nomis Foundation). Some of the graphical illustrations used in the manuscript were prepared using BioRender.com. We thank Core Facility Genomics (CF-GEN), Helmholtz Center Munich, Neuherberg, Germany for their support in running scRNAseq and spatial transcriptomics assays. We thank Mr. Marin Bralo at iBIO for his technical support throughout this study.

## Author contributions

Conceptualization and management of whole project,

H.S.B. and A.E.; methodology, H.S.B. formal analyses, L.S., L.K., H.S.B., M.A., Z.H.; investigation, H.S.B., C.M., S.J., D.J., M.I.T., S.P., H.E.P, J.F.T. I.S., K.R; visualization, L.H., H.S.B.; resources, A.R., B.T., M.B., F.J.T., S.U., H.A., O.B.; writing-original draft, H.S.B.; project administration, H.S.B.; writing – review & editing, H.S.B., A.E.; Funding acquisition, A.E.

## Declaration of interests

A.E. is co-founder of Deep Piction, GmbH. M.B. is a member of the Neuroimaging Committee of the EANM.

M.B. has received speaker honoraria from Roche, GE Healthcare, Iba, and Life Molecular Imaging; has advised Life Molecular Imaging, AC Immune, MIAC and

GE healthcare.

## Materials and Methods

### Animals

Mixed-gender C57BL/6 mice were obtained from Charles River Laboratories. The animals were housed under a 12-h light/dark cycle and had random access to food and water. The temperature was maintained at 18–23 °C, and the humidity was at 40–60%. The animal experiments were conducted according to institutional guidelines of the Klinikum der Universität München/ Ludwig Maximilian University of Munich and German mouse clinic (GMC) at Helmholtz Munich after approval of the Ethical Review Board of the Government of Upper Bavaria (Regierung von Oberbayern, Munich, Germany) and under European Directive 2010/63/EU for animal research. All data are reported according to the ARRIVE. Sample sizes were chosen based on prior experience with similar models.

### Cell culture

SB28-eGFP cells (44, 45) were provided by Walter-Brendel-Zentrum and cultured in DMEM containing MEM non-essential amino acids (1×), 1% penicillin-streptomycin solution (Thermo Fisher Scientific), and 10% fetal bovine serum (FBS, Biochrome). Cell cultures were maintained in the incubator at 37°C in a humidified and 5% CO2-conditioned atmosphere. Cells were passaged when the cell density in the flask reached 80% confluence. Cells were maintained in culture for up to 4 weeks after thawing, and cultures were passaged 5 to 10 times before utilization in animal experiments. Testing for mycoplasma infection was performed using MycoAlert PLUS Mycoplasma Detection Kit (Lonza Bioscience).

### Glioblastoma mouse model

C57BL/6 mice were purchased from Charles River (Sulzfeld, Germany) and acclimated for at least 1 week. At day 0, the mice were inoculated with 100,000 SB28-eGFP cells suspended in 2 μl of Dulbecco’s modified Eagle’s medium (DMEM) (Merck, Darmstadt, Germany) (glioblastoma mice, n = 7) or 2 μl of saline (sham mice, n = 4). For inoculation, mice were anesthetized with intraperitoneal injections of 10% ketamine (100 mg/kg) and 2% xylazine (10 mg/kg) in 0.9% NaCl. Anesthetized mice were immobilized and mounted onto a stereotactic head holder (David Kopf Instruments, Tujunga, CA, USA) in the flat-skull position. After surface disinfection, the skin of the skull was dissected with a scalpel blade. The skull was carefully drilled with a micromotor high-speed drill (Stoelting Co., Wood Dale, IL, USA) 2 mm posterior and 1 mm left of the bregma. By stereotactic injection, 100,000 cells in 2µl DMEM or 2µl of saline were applied with a 10-μl Hamilton syringe (Hamilton, Bonaduz, Switzerland) at a depth of 2 mm below the drill hole. Cells were slowly injected within 1 min, and after a settling period of another 2 min, the needle was removed in 1-mm steps per minute. After that, the wound was closed by suturing. Tumor and sham mice received [^18^F]FET small-animal PET recordings with an emission window of 0 to 60 min after injection of 16 ± 1 MBq [^18^F]FET using a Mediso PET/ computed tomography (CT) system (Mediso, Budapest, Hungary) between days 5 and 21 after inoculation. The last PET recording was scheduled close to perfusion and served for correlation with DISCO-seq. A CT was performed before the PET scan to allow attenuation correction. All small-animal PET experiments were conducted with isoflurane anesthesia (1.5% at the time of tracer injection and during imaging; delivery 3.5 liters/min). For all mice, a 40- to 60-minute frame was analyzed, and normalization of activity was performed by standardized uptake values (SUV).

### Protein conjugation with fluorophore and SARS-CoV2 spike S1 injection

Spike S1 protein (N501Y) was labeled with the fluorescent dye Alexa Fluor 647 according to the manufacturer’s protocol (Alexa Fluor 647 conjugation kit lightning link, Abcam, ab269823) as performed in our earlier study (11). Briefly, the candidate protein (250 μg) supplied by the manufacturer (SARS-CoV-2 (COVID-19) S1 protein (N501Y), His Tag (S1N-C52Hg, Acrobiosystems), were dissolved in 0.01 M PBS, pH 7.4. The modifying reagent was added to the protein solution. The solution was then mixed gently and transferred to a lyophilized Alexa Fluor 647 material. After 15 minutes of incubation at room temperature, the reaction was terminated by adding the quencher reagent. The fluorophore and the S1 protein were mixed in 1:1 ratio and had a final concentration of 1 mg/ml and injected via tail vein at the concentration of 10 μg/mouse for 30 minutes.

### Perfusion, fixation and tissue preparation

Mice were deeply anesthetized using a combination of ketamine and xylazine (concentration, 2 ml per 1 kg i.p.). Afterward, the chest cavity of mice was opened for intracardial perfusion with cold PBS using 100–125 mmHg pressure for 5 min at room temperature until the blood was washed out.

For the fresh frozen condition, tissues were rapidly dissected immediately following perfusion with ice-cold phosphate-buffered saline (PBS, pH 7.4) to remove blood contaminants. Samples were carefully blotted to remove surface moisture, snap-frozen in liquid nitrogen, and stored at −80 °C until use. All instruments and surfaces were RNase-free, and handling was performed swiftly to minimize RNA degradation. This approach preserved native RNA content without any fixation or chemical alteration, serving as the baseline for subsequent chemical preservation and clearing optimizations. For fixations, tissues were perfused with ice-cold PBS followed by freshly prepared 4 % paraformaldehyde (PFA) or 3 % glyoxal or RNAlater. Fixed tissues were post-fixed by immersion in the respective fixative solution for 3-4 hours at 4 °C to ensure uniform penetration. Following fixation, tissues were washed thoroughly in PBS to remove residual fixative and equilibrated either in RNAlater at 4 °C for 24 hours to stabilize RNA or proceeded with diverse clearing protocols.

### Optimized DISCO-seq chemistry

The optimized DISCO-seq workflow was developed to combine efficient tissue clearing, whole organ imaging with preservation of high-quality RNA for downstream transcriptomics. The process began with transcardial perfusion of mice (mouse on ice bed while perfusing) using ice-cold PBS, followed by perfusion of ice cold 4 % PFA prepared in nuclease free PBS and post-fixed in 4 % PFA at 4 °C for 3-4 hours. Fixed tissues were washed 2-3 times quickly in cold PBS and proceeded to tetrahydrofuran (THF, Roth, CP82.1)/dichloromethane (DCM, Sigma, 360,538) -based clearing. Dehydration was performed through a graded series of freshly prepared, anhydrous THF) solutions (50 %, 70 %, 90 % and x2 100 %, v/v in nuclease-free water), each for 3 hours at at 4 °C with gentle agitation. Residual water was removed by two additional incubations in 100 % THF. Samples were then immersed in DCM for 20–30 minutes as an intermediate solvent step to enhance clearing efficiency, followed by immersion in BABB (1:2 benzyl alcohol:benzyl benzoate, Sigma, 24,122 and W213802) until full transparency was achieved.

For tert-butanol based clearing, a gradient of tert-butanol in distilled water (50 %, 70 %, 90 % and x2 100%, v/v in nuclease-free water), each for 3 hours with gentle agitation at 32-34°C, followed by immersion in DCM for 20-30 min at room temperature and finally incubated with the refractive index matching solution BABB-D15 containing 15 parts BABB, 1 part diphenyl ether (DPE) (Alfa Aesar, and 0.4% Vol vitamin E (DL-alpha-tocopherol, Alfa Aesar, A17039), for at least 6 h at room temperature until achieving transparency.

To minimize RNA degradation during hydrophobic clearing, all solvent exchanges were performed rapidly, under anhydrous and RNase-free conditions. The final cleared samples were either immediately proceeded for optimized rehydration protocol or imaged using light sheet microscope in refractive-index matchimg solution (BABB). The integration of cold perfusions, optimized fixation, optimized solvent choice, incubation timings/ temperature, and handling defined the DISCO-seq chemistry, which preserves both structural integrity for imaging and RNA quality for sequencing.

### Optimization of DISCO cleared tissue prior to RNA extraction, cryosectioning and probe hybridization

The course of tissue clearing and imaging in BABB makes the tissue brittle and hard to process further. To solve this, we optimsed reverse clearing of samples with the respective clearing solutions to be able to process for downstream steps. For the DISCO-seq cleared samples, the samples were sequentially rehydrated using reverse gradients of THF (100 %, 90 %, 70 %, 50% v/v in nuclease-free water), each for 90 minutes. Particularly, the water based steps (i.e., 90 %, 70 %, 50%) were performed at 4 °C with gentle agitation. Samples were then washed twice with cold PBS at 4 °C for 10 minutes each. Samples were then either cryopreserved with 30% sucrose solution in 4°C or directly processed for RNA extractions or processed for flex assay protocol.

### DISCO clearing of whole mouse bodies

We performed tissue clearing based on the DISCO protocol for whole mice as previously described (11) adapted to DISCO-seq chemistry. For the tissue clearing of mice after intravenous injection of spike S1, perfusions/ fixations were performed as described above in DIS-CO-seq protocol. After that mouse bodies were placed in a 300 ml glass chamber and incubated in 180-200 ml of the following gradient of THF in distilled water in a fume hood with gentle shaking: (50 %, 70 %, 90 % and x2 100 %, v/v in nuclease-free water), 6 h for each step, followed by 2 h in DCM, and finally in BABB solution until optical transparency. Rehydrations of organ/region of interest were processed as described above.

### Light sheet fluorescence imaging

Single plane illuminated (light-sheet) image stacks were acquired using an UltraMicroscope Blaze (Miltenyi Biotec), featuring an axial resolution of 4 μm with following filter sets: ex 470/40 nm, em 535/50 nm; ex 545/25 nm, em 605/70 nm; ex 640/40 nm, em 690/50 nm. Whole brains were imaged individually using high magnification objectives: 4× objective (4× corrected/0.28 NA [WD = 10 mm]). High magnification tile scans were acquired using 20-35% overlap and the light-sheet width was reduced to obtain maximum illumination in the field. Processing, data analysis, 3D rendering and video generation for the rest of the data were done on an HP workstation Z840, with 8 core Xeon processor, 196 GB RAM, and Nvidia Quadro k5000 graphics card and HP workstation Z840 dual Xeon 256 GB DDR4 RAM, nVidia Quadro M5000 8GB graphic card. We used Imaris (Bitplane), Fiji (ImageJ2), Vision 4D (Arivis) and syGlass (for 3D and 2D image visualization). Tile scans were stitched by Fiji’s stitching plugin49.

### Optimizations of DISCO cleared tissue and nuclei extraction

#### Nuclei isolation from DISCO cleared tissue

Working in a fume hood, use a 1 mm punch biopsy tool to take a sample from the DISCO cleared mouse brain in a petri dish holding it with forceps. Place the punches in a safe lock 1.5 mL Eppendorf tube. Wash the punches with a series of 1mL ethanol washes to rehydrate the tissue. Incubate the tissues for 2 mins (2×100% EtOH, 2x 75% EtOH, 2x 50% EtOH). Incubate at 80°C for the first three washes to begin reverse cross-linking the tissue for a total of 6 mins. The remaining washes can be completed at room temp. After rehydration is complete work on ice and add 1 mL of 1X ST buffer: [146 mM NaCL; 21 mM MgCl2; 1 mM CaCl2; 10 mM Tris-HCL; 40 U/mL RNase Inhibitor; 0.01% BSA] to remove residual ethanol. Remove ST and add 500 µl of CST lysis buffer: [146 mM NaCL; 21 mM MgCl2; 1 mM CaCl2; 10 mM Tris-HCL; 0.5% CHAPS; 40 U/ mL RNase Inhibitor; 0.01% BSA] and then chop for 10 mins using Noyes Spring Scissors - Tungsten Carbide/ Straight (Fine Science Tools 15514-12) and then using Douncer (Sigma D8938). Filter through a prewetted 30 µm nylon-mesh filter (CellTrics 04-004-2326), using 2.5 mL of ST buffer to both prewet the filter and rinse out the tube. Add 2 mL of ST buffer to the filter to rinse off any residual nuclei into collection for a total of 5 mL of nuclei. Centrifuge in a 4 °C swing bucket centrifuge for 5 mins at 500g, slow the break speed to avoid the pellet coming loose. Remove supernatant and resuspend in 50 µL of ST buffer per initial number of punches taken. Prior to resuspension, a 10 µL aliquot of the nuclei pellet can be used for RNA extraction and QC using the FormaPure FFPE RNA extraction kit (Beckman Coulter C19157). Count nuclei using DAPI and a hemocytometer or automated cell counter such as Cellometer with the AO/PI stain. Add Ruby Vybrant™ DyeCycle™ Ruby Stain, (ThermoFisher #V10309) to sample 1:500 (2 µL for 1mL) or DAPI (1:5000) to the nuclei suspension. Sorted using either the Sony SH800 or BD Aria fusion to wells in the 96 well PCR plate containing 5 µL of Qiagen TCL lysis buffer (1031576) with 1% BME. Nuclei were sorted into a gradient where we used 500 nuclei, then 100, 50, 10, and single. Nuclei from DISCO-seq were sorted into separate plates from nuclei from tert-butanol.

### Modifications to the SS2 protocol for DISCO sorted nuclei

We added a decrosslinking step to the published protocol adding Proteinase K (NEB P8107S) at a final concentration of 0.2 µg/µL, adding 1ul of volume to each well and incubating at 55 °C for 15 mins and 80 °C for 15 mins in a thermal cycler. After decrosslinking is complete, we added a positive control well with 1 ng of mouse brain RNA integrity score of 9. If added beforehand, this decrosslinking treatment would degrade the RNA. We then proceeded to process the plate through the SS2 protocol (28). DISCO-seq and tert-butanol plates were processed side by side until the SPRI elution step where the samples were then transferred into the same plate for reverse transcription and library construction.

### Bulk nuclei and single nucleus RNA sequencing

SnRNA sequencing was performed on the Illumina NextSeq500 using the High Output Kit v2 (75 cycles) FC-404-2005 with the read structure as follows: Index 1 - 8bp, Read 1 - 38bp, Read 2- 38bp, Index 2 - 8bp.

### Fixed RNA profiling of mouse brain and small intestine tissue using 10x Genomics flex assay

#### Single-cell dissociation and fixation compatibility

Frozen tissue was thawed on ice and enzymatically dissociated using the 10x Genomics Fixed RNA Profiling (FRP) protocol, which is optimized for PFA-fixed tissues. Tissue chunks were incubated in dissociation enzymes provided in the FRP kit and triturated gently to obtain a single-cell suspension. The resulting suspension was passed through a 30 µm cell strainer to remove aggregates. Cell concentration and viability were measured using a hemocytometer and trypan blue exclusion.

#### Probe Hybridization and Sample Barcoding

The fixed and permeabilized cells were processed using the 10x Genomics Chromium Flex Gene Expression assay (singleplex 4-sample kit was used for fresh frozen vs DISCO-seq cleared sample comparison; multiplex – 4rxns x 4BC - sample kit was used for mouse glioblastoma and spike S1 protein studies) following the manufacturer’s protocol. Approximately 8,000– 10,000 cells per sample were targeted for capture. Cells were incubated with a pool of target-specific barcoded probes covering ∼19,000 mouse genes (Chromium Mouse Probe Set v1.0.1), allowing hybridization to target mRNA sequences without the need for reverse transcription at this stage. After probe binding, unbound probes were removed through a series of washes.

#### Gel Bead-in-Emulsion (GEM) Generation and Library Construction

Probe-labeled cells were loaded onto a Chromium Controller X (10x Genomics), where individual cells were encapsulated into gel bead-in-emulsions (GEMs) along with reagents for gap-filling and ligation. Unique molecular identifiers (UMIs), sample barcodes, and cell barcodes were incorporated during GEM formation. Following amplification and clean-up steps, indexed libraries were generated according to the Flex assay specifications.

### Sequencing

Final libraries were sequenced on Illumina NovaSeq 6000 & NovaSeq X Plus sequencers using paired-end sequencing parameters optimized for the Flex assay (e.g., Read 1: 28 bp, Read 2: 90 bp, Index 1: 10 bp, Index 2: 10 bp). The target sequencing depth was approximately 20,000 reads per cell.

### Quality control and preprocessing

Count data was generated with Cell Ranger (46) (v8.0.0) using the mm10-2020-A reference transcriptome for the fresh frozen versus cleared brain samples analysis and Cell Ranger (v9.0.1) with the GRCm39-2024-A reference transcriptome for the glioblastoma and Spike S1 analysis. The count matrices were analyzed using Python (Scanpy v1.11.2, and v1.11.3, 47) and R. Ambient RNA contamination was removed using SoupX (v1.6.2, 48). Doublets were identified using scrublet (47) with default parameters and removed from the dataset. Additional low-quality cells were removed based on three quality metrics: (1) number of expressed genes in a cell (< 300), (2) total UMI counts (< 400), and (3) mitochondrial UMI counts (> 15% for fresh frozen brain samples, 7% for cleared brain samples, and 4.5% for cleared small intestine samples). Count data was normalized using scanpy.pp.normalize_total and a log1p transformation was performed. 6000 highly variable genes were computed with the sample name as the batch key. For the fresh frozen versus cleared brain samples analysis, batch correction was performed using scVI (scvi-tools v1.3.3, 36, 37) with n_layers=2, n_latent=30, and gene_likelihood=“nb”.

### Cell type annotation

Principal components, nearest neighbors, and a UMAP were generated and leiden clustering was performed with a resolution of 1.5. For the brain samples, automated cell type annotation using MapMyCells (35) (v1.6.1, RRID:SCR_024672) from the Allen Brain Institute was performed locally and cells with a sub-class bootstrapping probability of less than 0.5 were removed. For the gut samples, manual annotation was performed per cluster using a list of marker genes (and by studying the differentially expressed genes for certain clusters). Subclustering with the default resolution of 1 was also performed for more detailed annotations. Cells co-expressing marker genes of different cell types or for which the annotation is unclear were removed from the dataset. Cells due to contamination, including adipocytes and pancreatic acinar cells, were also removed.

### Differential gene expression analysis and gene ontology enrichment

Differential gene expression analysis was performed using scanpy.tl.rank_genes_groups with method=‘wilcoxon’ for each cell type (and region or condition) separately. Cell types for which less than 50 cells were available (per region or condition) were not studied. The threshold for the FDR adjusted p-values was chosen to be 0.01. If the number of up or downregulated genes was more than 10, gene ontology enrichment was performed using GProfiler (48) provided with a maximum of 100 up or downregulated genes.

### Co-registration of PET/CT with lighsheet microscopy data

The CT and PET scans of the mouse head were acquired as co-registered volumes. The skull was manually segmented from the CT images, and the interior of the skull was used as the brain mask. This brain mask was as the moving volume in the registration process, which was aligned to the light sheet scan (fixed volume). Registration was performed through a series of rigid, affine, and B-spline transformations using elastix (49) (v5.01), with 3-6 levels of resolution pyramids. The resulting transformation parameters were then applied to the PET volume using nearest-neighbor interpolation, aligning it with the light sheet scan. In the second registration, the average template from the Allen Brain Atlas (50) (CCFv3) was used as the moving volume, and the same registration process was applied. Finally, the transformation parameters were re-applied to the atlas annotation volume, again using nearest-neighbor interpolation, ensuring that the atlas annotations were correctly aligned with the light sheet scan.

### High-resolution spatial transcriptomics with 10x Genomics Xenium assay on cleared mouse glioblastoma brain Sections

#### Tissue Clearing and Sectioning

Adult mouse brains were fixed in 4% paraformaldehyde (PFA) for 3 hrs at 4 °C and processed using the DISCO clearing protocol to enhance tissue transparency while preserving RNA integrity. Clearing was achieved through stepwise dehydration in tetrahydrofuran (THF) followed by benzeyl alcohol: benzyl benzoate (1:2) (BABB) immersion for refractive index matching. Tissues were then rehydrated gradually and embedded in OCT prior to cryosectioning. Cryosections were cut at 12 µm thickness using a cryostat and mounted onto Xenium-compatible glass slides. Sections were stored at –80 °C until further processing.

#### Xenium In Situ Spatial Transcriptomics Assay

Spatial transcriptomic profiling was carried out using the Xenium Analyzer (10x Genomics) following the manufacturer’s protocols. We used the Xenium Mouse Gene Expression Panel v1.0, which targets ∼247 genes selected to capture key mouse brain cell types and functional states. Following tissue permeabilization, multiplexed, barcoded RNA probes were hybridized directly to mRNA molecules within the tissue section. In situ signal amplification and iterative imaging were performed using cyclic fluorescent labeling chemistry, enabling spatial decoding of each transcript at subcellular resolution without cDNA synthesis.

#### Imaging and Cell Segmentation

High-resolution imaging was performed on the Xenium Analyzer, capturing fluorescent signals across multiple imaging rounds. DAPI staining was used to identify nuclei, and segmentation was performed using the Xenium Onboard Analysis pipeline, which includes both nuclear expansion-based segmentation and cell membrane-based segmentation (using spatial transcript distribution and image features). This dual segmentation approach allowed more accurate delineation of cell boundaries and spatial localization of transcripts.

#### Data Processing and Output

Spatial transcriptomic data were processed using Xenium Onboard Analysis (XOA), generating:

- Cell-by-gene expression matrices
- Spatial coordinates for each transcript and nucleus
- Segmented cell masks (nucleus-expanded and membrane-defined)
- Per-cell and per-transcript quality metrics

Output files were visualised and explored using Xenium Explorer and the spatial data python package. Subsequently, Scanpy was used for downstream analysis and comparison with scRNAseq data.

### Comparison of DISCO-seq and Spatial Xenium Datasets

Single-cell DISCO-seq and spatial Xenium transcriptomic datasets were generated from matched glioblastoma tissue samples. Raw count matrices were processed using Scanpy (v1.9.6) and normalized to the median count per cell followed by log-transformation. Cell-type identities were assigned using the mapmycells framework with the 10x whole mouse brain CCN20230722 reference. Only cell assignments with a mapping confidence score ≥ 0.8 were retained. Cell types with fewer than 80 cells in either dataset were excluded from further analysis to ensure robust representation. Marker genes were selected based on a combination of curated canonical markers and the top differentially expressed genes identified by the Wilcoxon rank-sum test implemented in Scanpy. Genes were further filtered for detection in both modalities to enable direct comparison.

## Supplementary video legends

**Video S1**

Whole-Brain 3D Imaging of a Glioblastoma Mouse Model. Showing large primary tumor, few discrete aggregated tumors and their trafficking via corpus callosum. SB28-eGFP tumor cells are in white, annotated metastatic tumors are in green, autofluorescence is in gold.

**Video S2**

3D reconstruction of whole mouse body showing the localization of spike S1 protein. Zooming into parts of duodenum and ileum. Autofluorescence is in gold, Spike S1 is in cyan.

---

**Figure S1.**
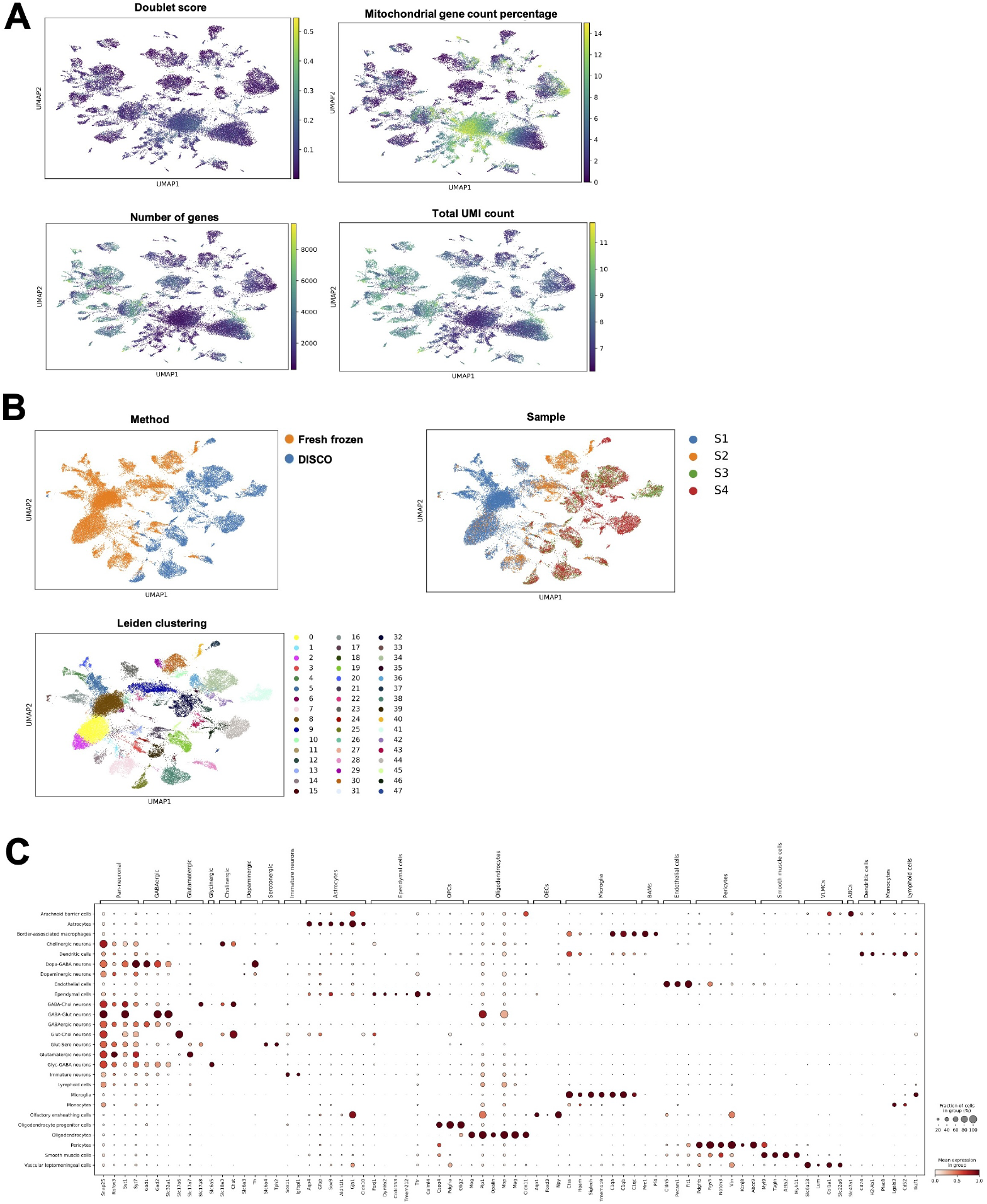
Comparison of single cell RNA sequencing data in DISCO cleared vs. fresh frozen mouse brain samples – quality control. **(A)** Quality control of processed cells after integrated scVI embedding and cell annotations by looking at the fraction of doublet cells, percentage of mitochondrial gene counts mapped on the cell clusters, number of genes, and number of UMI counts (log1p). **(B)** UMAP by condition (upper left plot), by sample (right plot) and Leiden clustering of all samples before cell annotations (bottom plot). **(C)** Dot plot of canonical cell-type marker genes showing the mean expression and fraction of expressing cells used for cell annotations.

**Figure S2.**
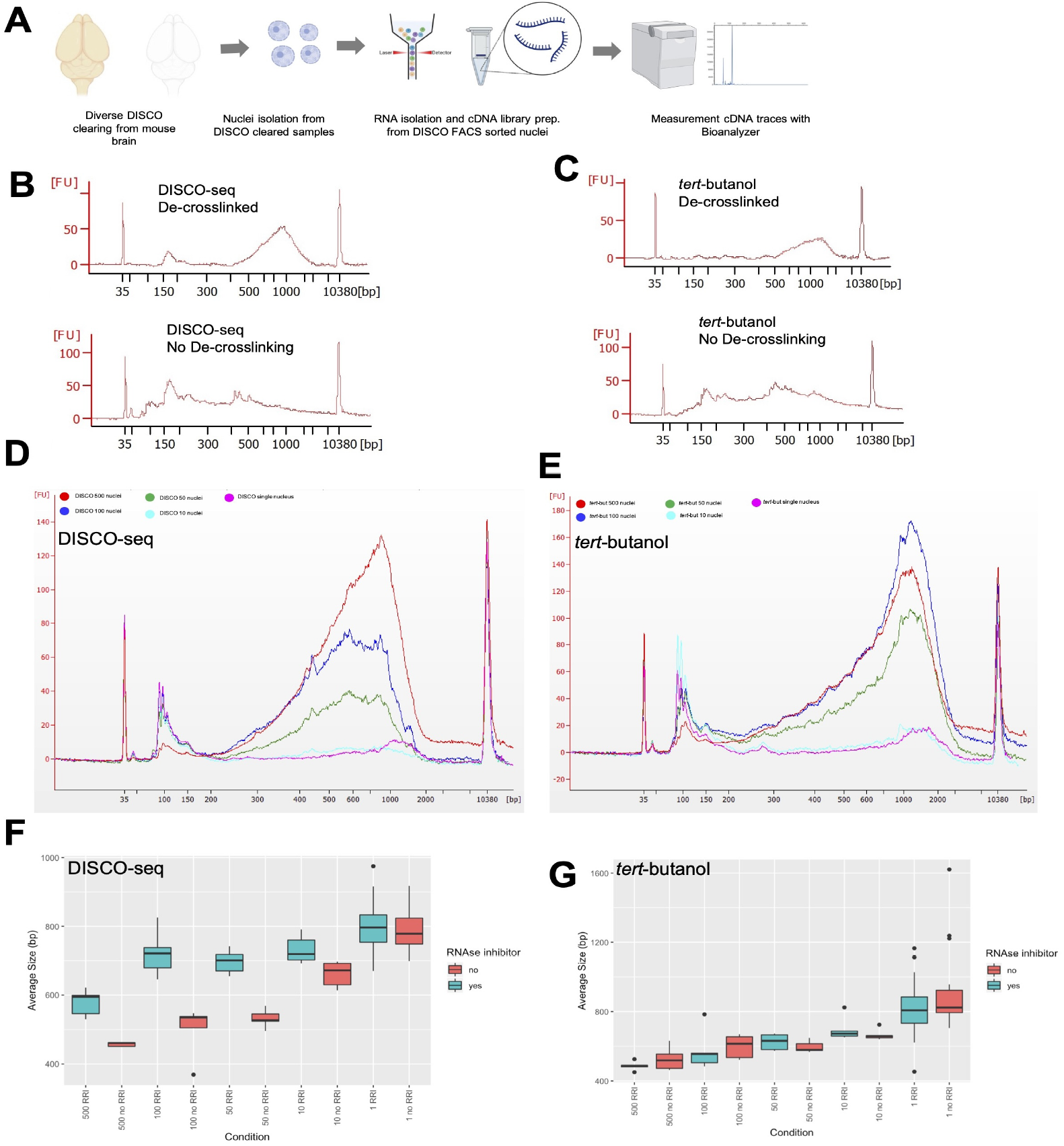
Improvement of cDNA library preparation of diverse DISCO cleared nuclei suspension. **(A)** Schematic of DISCO clearing, isolation of RNA from bulk tissue and DAPI sorted nuclei suspension, subsequent RNA extraction and cDNA library preparation. **(B)** Electropherograms showing the cDNA traces from DISCO treated nuclei indicating the effect of de-crosslinking prior to SMART-seq2 (SS2). **(C)** Electropherograms showing the cDNA traces from tert-butanol treated nuclei indicating the effect of de-crosslinking prior to SMART-seq2 (SS2). (D) Average cDNA library size from DISCO cleared nuclei suspension from 500, 100, 50, 10 and single nucleus. **(E)** Average cDNA library size from tert-buanol cleared nuclei suspension from 500, 100, 50, 10 and single nucleus. **(F)** Effect of RNase inhibitor on the average cDNA library size from DISCO cleared nuclei suspension from 500, 100, 50, 10 and single nucleus. **(G)** Effect of RNase inhibitor on the average cDNA library size from tert-butanol cleared nuclei suspension from 500, 100, 50, 10 and single nucleus.

**Figure S3.**
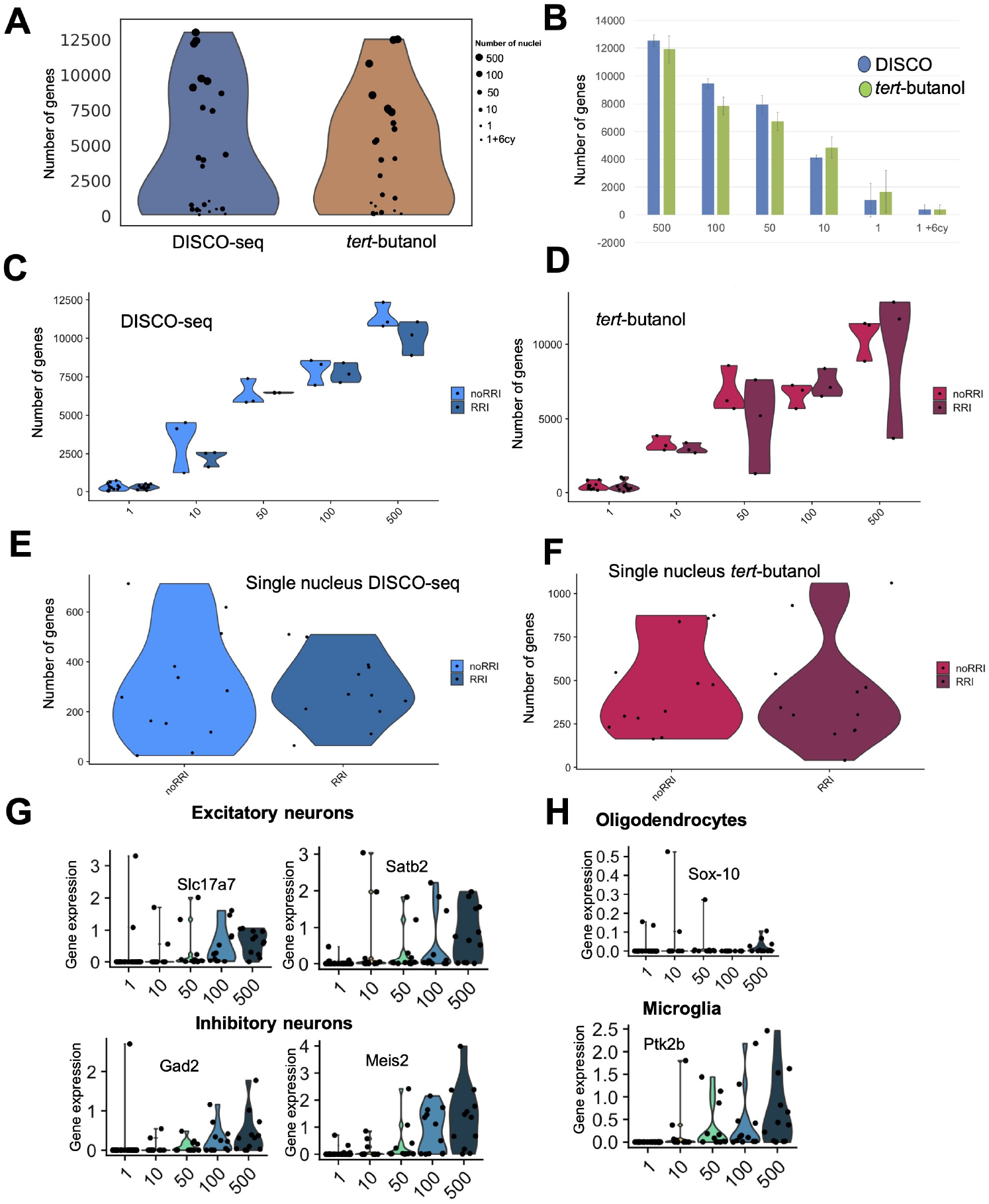
Bulk and single nucleus RNA sequencing (snRNAseq) from DISCO cleared mouse brain samples. **(A)** RNA Sequencing results showing the number of detected genes across diverse number of isolated nuclei from DISCO-seq vs. tert-butanol cleared samples. The size of dots in violin plot relate to the number of nuclei in the SMART-seq2 reaction. 3 replicates for the bulk samples (500, 100, 50, 10) and 6 replicates for the single nucleus were used. **(B)** Same data plotted as bar graph to show the variation in the detected number of genes as we go from bulk sample towards single nucleus and effect of additional 6 cycles in PCR (1+6cy) sample. **(C)** Effect of RNase inhibitor (RRI) on the detection of number of genes in DISCO cleared samples. **(D)** in tert-butanol cleared samples. **(E)** Detected genes in single nucleus in DISCO cleared samples showing >200 genes detected for 17 out of 24 nuclei (not separated based on RNase treatment). **(F)** Detected genes in single nucleus in tert-butanol cleared samples showing >200 genes detected for 20 out of 24 nuclei (not separated based on RNase treatment). **(G)** Exemplary marker genes associated with neurons (excitatory, upper panel, inhibitory neurons, lower panel) **(H)** glial cells (oligodendrocytes, upper panel, microglia, lower panel) with their expression profile from DISCO cleared samples.

**Figure S4.**
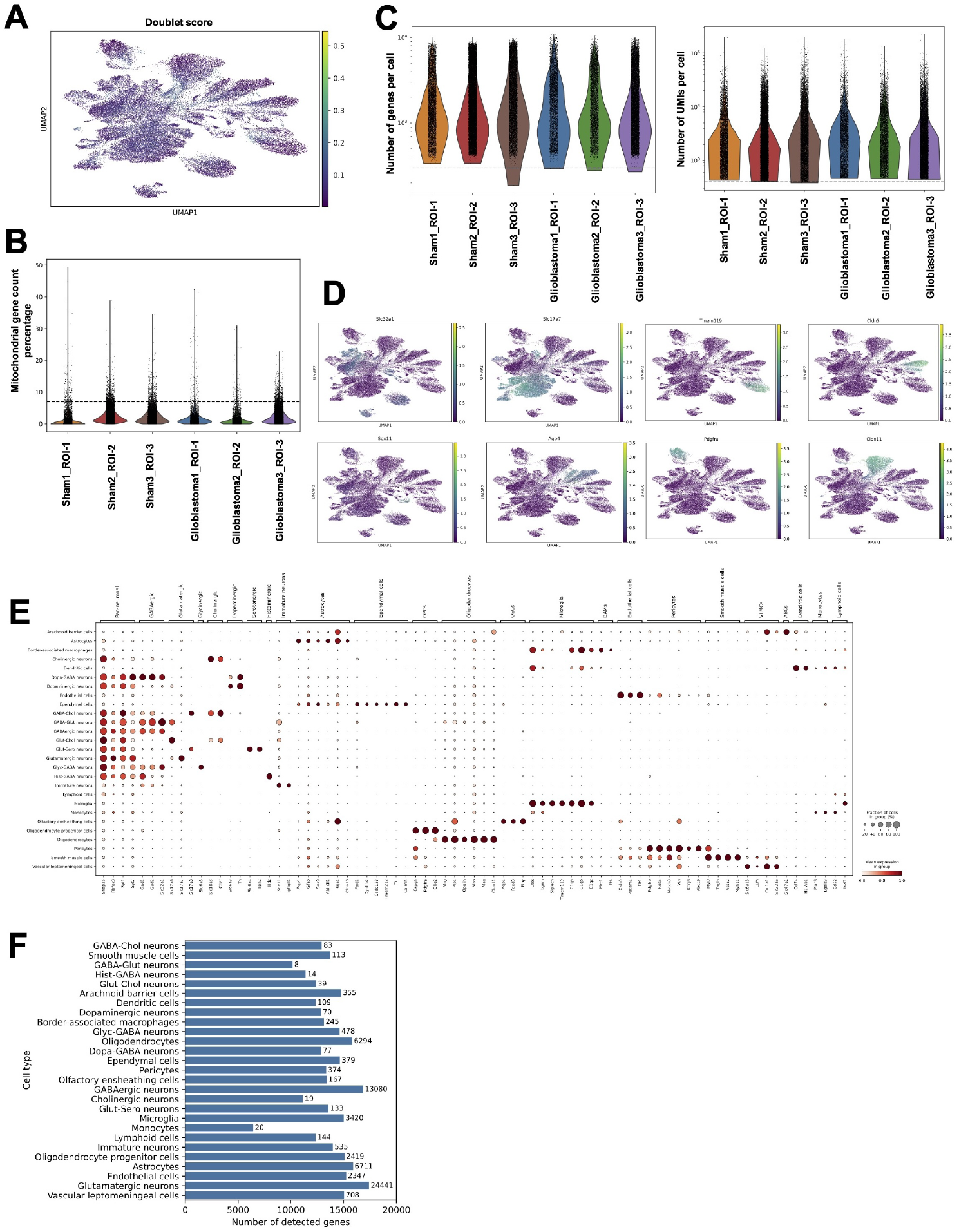
DISCO-seq preserve the expected cell type and cell proportion across glioblastoma and sham brains. **(A)** Showing doublet score across conditions and cell types as proxy of quality of processed cells. **(B)** Percentage of mitochondrial reads across sham and DISCO cleared samples. **(C)** Number of genes recovered per cell in each condition (left panel), number of UMIs per cell in each condition (right panel). **(D)** UMAP showing distribution of various cell types in mouse brain according to their canonical marker genes. **(E)** Dot plot of cell-type marker genes showing the mean expression and fraction of expressing cells used for cell annotations. **(F)** Bar graph showing number of detected genes per annotated cell type, numbers on top of the each bar indicates the total cell number for each cell type.

**Figure S5.**
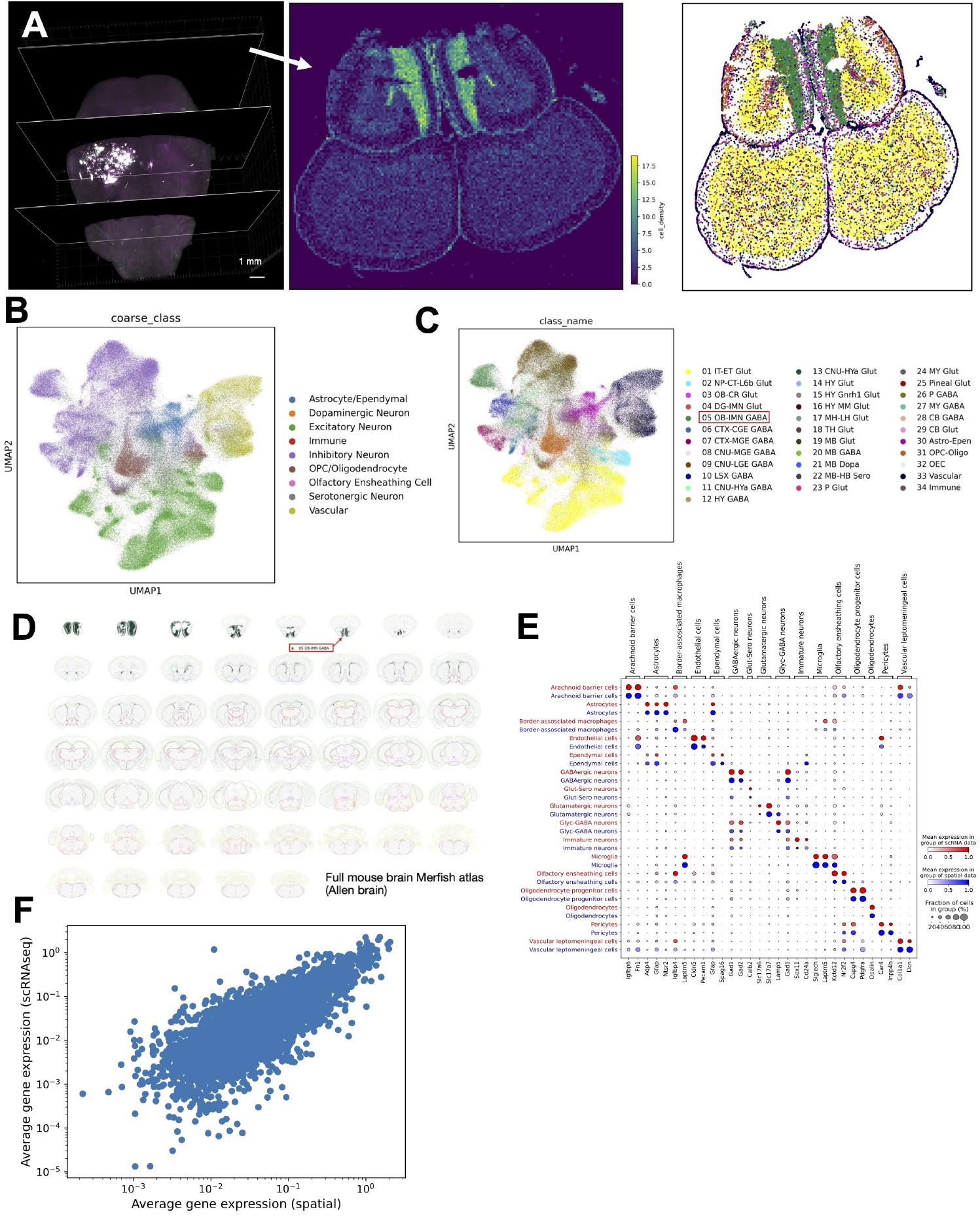
Targeted spatial transcriptomics using Xenium mouse brain panel on DISCO cleared glioblastoma brains. **(A)** 3D whole brain DISCO-seq clearing and imaging guided slice selection for single cell spatial transcriptomics, showing cell density and cell distribution maps on the distant ROI section. **(B)** Coarse cell annotations using Mapmycells algorithm for cell type mapping showing diverse cellular identities. **(C-D)** Fine cell type annotations to show further intact sub-types of cellular population identified after DISCO clearing and showing exemplary regional specificity of cell-type class of GABAergic OB immature neuron (OB-IMN-GABA) type which align well with MERFISH brain atlas. **(E)** Dot-plot showing mean normalized gene expression (color intensity) and fraction of expressing cells (dot size) in matched cell-type groups between DISCO-seq data (in red) and spatial xenium data (in blue). **(F)** Scatterplots of average gene expression (Pearson correlation, R=0.81) between DISCO-seq glioblastoma data vs. Spatial xenium glioblastoma data.

**Figure S6.**
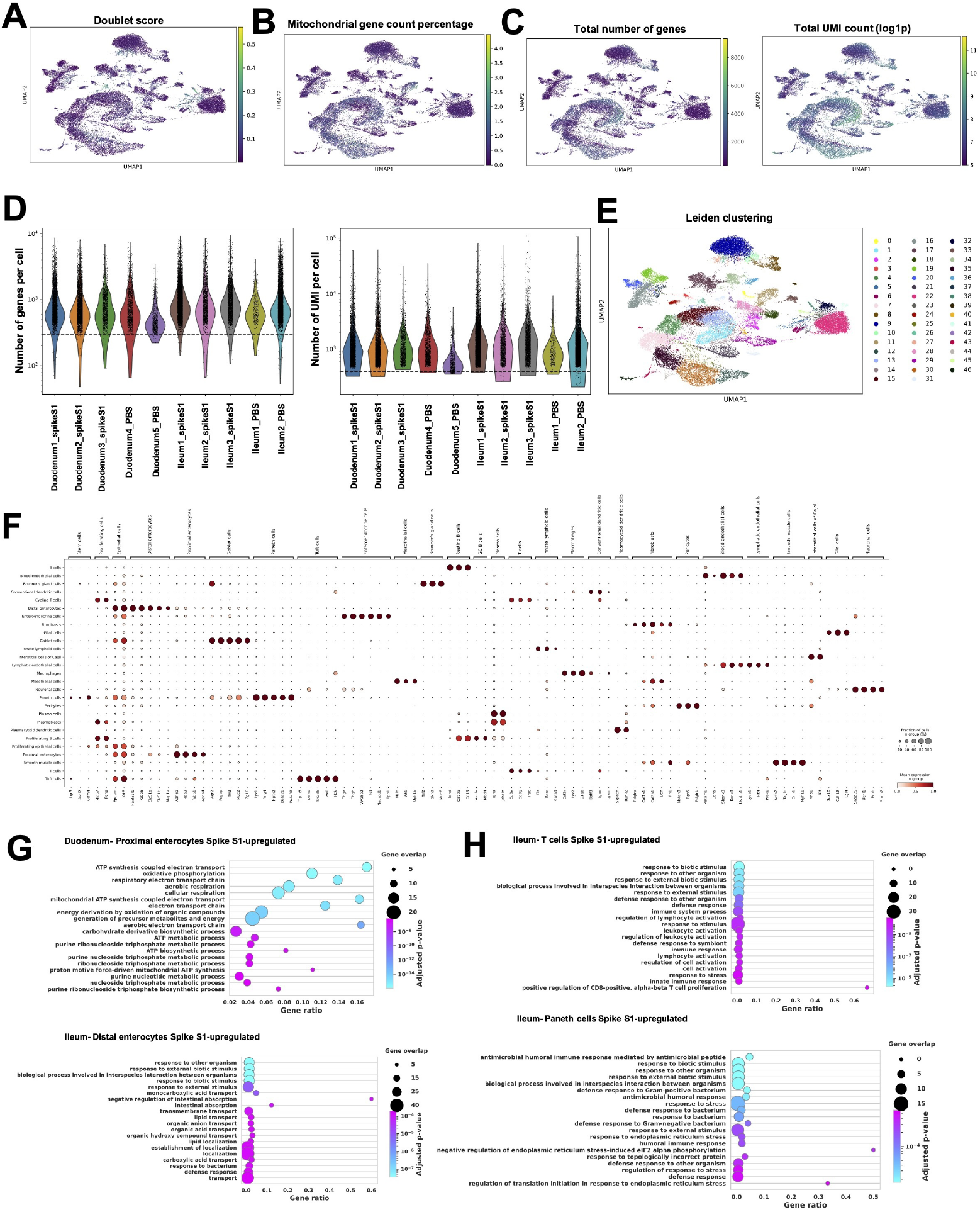
Spatial molecular analyses in sub-regions of small intestine using DISCO-seq. **(A)** Showing doublet score across conditions and cell types as proxy of quality of small intestine processed cells. **(B)** Percentage of mitochondrial reads across samples and cell types **(C)** Number of genes (left panel) and UMIs (right panel) recovered per cell in each condition mapped on the cell clusters. **(D)** Number of genes and counts per sample type. **(E)** Leiden clustering of all samples before manual cell annotations. **(F)** Dot plot of canonical cell-type marker genes showing the mean expression and fraction of expressing cells used for cell annotations. **(G)** Upregulated Gene ontology (GO) terms in enterocytes from Duodenum (upper left panel) and ileum (lower left panel) after spike S1 injection. **(H)** Upregulated GO terms in T cells (upper right panel) and Paneth cells (bottom right panel) in ileum after S1 spike injection.

